# Exploring the Role of APC in Modulating Chemotherapeutic Response in Triple-Negative Breast Cancer cells

**DOI:** 10.64898/2025.12.31.697243

**Authors:** T. Murlidharan Nair, Monica K. VanKlompenberg, Jenifer R. Prosperi

## Abstract

Triple-negative breast cancer (TNBC) lacks targeted therapies and often develops chemoresistance. Since loss of the tumor suppressor APC is common in TNBC, we investigated how APC depletion alters transcriptional adaptation to chemotherapy using RNA-seq profiling of MDA-MB-157 cells and APC knockdown derivatives under control, cisplatin, and paclitaxel treatment. Pathway analyses of the transcriptomes showed that APC knockdown induces extensive remodeling of DNA-damage response, mitochondrial metabolism, inflammatory signaling, and chromatin accessibility, establishing stress-adapted transcriptional states aligned with paclitaxel tolerance (APC_shRNA1) or cisplatin resilience (APC_shRNA2). Machine-learning feature selection (Random Forest + PLS-DA) identified a 43-gene discriminant signature enriched for regulators of cell cycle and DNA repair (CCNB3, ORC1, E2F2, UNG), cytokine signaling (CXCL2, IL11), and metabolic support. These findings suggest that APC loss primes TNBC cells for chemotherapy persistence through an energetically reinforced, transcriptionally flexible survival program, highlighting DDR–OXPHOS–translation and inflammatory circuits as potential therapeutic vulnerabilities.

## 1. Introduction

Breast cancer is a heterogeneous disease and one of the leading causes of mortality in women worldwide [1, 2]. Clinically it is categorized into three therapeutic groups: the estrogen receptor (ER) positive group [3], the HER2/ERBB2/Neu amplified group [4], and triple-negative breast cancers (TNBCs) [5, 6] that lack receptors for estrogen, progesterone and HER2. TNBC is a high-grade cancer characterized by the expression of proliferation markers. Molecular signatures associated with the disease have been utilized to determine their prognostic value [3]. TNBCs have poorer prognosis compared to ER+ and HER2+ breast cancers, and they tend to metastasize to visceral organs and the central nervous system. Gene expression studies of breast cancer have revealed that they tend to self-segregate in different categories known as Luminal A, Luminal B, HER2 amplified and basal-like. The basal-like category predominantly consisted of TNBCs, suggesting that TNBCs may represent a molecularly defined category [7]. However, TNBCs have been shown to be heterogeneous; while a significant proportion are basal-like, they can also exhibit Luminal A, Luminal B, HER2, or Normal-like characteristics and have been further molecularly characterized [8]. Therefore, identifying drug targets based on molecular genomic signatures has proven challenging [9], and response to therapy varies depending on molecular subtype signatures [10–13]. Consequently, the therapeutic options also vary due to the heterogeneous nature of the disease [14–16].

Studies on the association between the tumor suppressor gene, *Adenomatous Polyposis Coli* (APC), and breast cancer have revealed that certain genetic variants [17] and/or epigenetically silenced versions of the gene are capable of activating Wnt/β-catenin pathway [18–21] to regulate tumor development. Furthermore, there is evidence to suggest that heterozygosity of APC could also enhance tumor development [21]. This underscores the critical role of APC in maintaining normal cellular state through both Wnt-dependent and Wnt-independent mechanisms. In addition to its involvement in Wnt/β-catenin signaling, APC plays a crucial role in regulating the mitotic spindle, which facilitates proper chromosome segregation [22–24], cell cycle control [25] and apoptosis [26]. APC has also been demonstrated to localize to the cell border and regulate cell motility [27].

Treatment of cancers using chemotherapy may be either neoadjuvant or adjuvant, and often involve a combination of drugs [28–32]. A common mechanism for the acquisition of resistance results from the overexpression of one or more energy-dependent transporters that are capable of effluxing anticancer drugs from the cell [33]. Other mechanisms for chemotherapeutic resistance include epigenetic modifications that alter key posttranslational modifications [34, 35]. Additionally, chemotherapeutic resistance has been linked to Wnt signaling [36, 37] and APC has been associated with mediating chemotherapeutic resistance independent of Wnt signaling [38, 39].

Understanding the role of APC in chemotherapeutic resistance is therefore critical for the development of novel therapeutic strategies for breast cancer. We previously demonstrated that knockdown of APC in the human TNBC cell line MDA-MB-157 (APC shRNA1 and APC shRNA2) results in differential resistance to paclitaxel or cisplatin, respectively [38]. To further investigate the molecular basis of this resistance, we performed transcriptomic analysis of MDA-MB-157 and the two APC knockdown lines, both with and without cisplatin and paclitaxel treatment. To define the transcriptional adaptations underlying APC-dependent drug response, we performed pathway enrichment analysis using GOAT (Gene Ontology & Association Tool) [40] across Hallmark Gene Sets, GO ontologies (BP, MF, CC), and Oncogenic Signatures, enabling systematic characterization of cellular rewiring associated with chemotherapeutic stress and resistance. Further, a combination of machine learning methods was employed to delineate discriminant signatures.

## 2. Methods

### 2.1 Cell Culture

The human TNBC cells, MDA-MB-157 (ATCC, Manassas, VA) were cultured in RPMI 1640 media supplemented with 10% fetal bovine serum, 1% penicillin/streptomycin, 25 mM HEPES, and 1:5000 plasmocin. Authentication of MDA-MB-157 cells was performed using STR DNA profiling by Genetica DNA Laboratories (Burlington, NC). Routine passaging of all cells was carried out using 0.25% trypsin/EDTA, and they were maintained at 37°C with 5% CO2. Transduced MDA-MB-157 cells were cultured in media containing 1.5 µg/mL puromycin (Sigma-Aldrich).

### 2.2 Drug Treatment

Cells were treated for a 24-hour period with either paclitaxel (0.078 µM), cisplatin (48 µM), or a solvent negative control. RNA was isolated from MDA-MB-157 and APC shRNA cells using Tri Reagent (Molecular Research Center, Cincinnati, OH), and cDNA synthesis was performed using iScript (Bio-Rad Laboratories, Hercules, CA). RNA sequencing was conducted to identify changes occurring in APC shRNA cells compared to MDA-MB-157 cells. Samples from control-treated and chemotherapy-treated cells were analysed through the Genomics Core Facility at the University of Notre Dame.

### 2.3 Expression profiling

Total RNA concentration and integrity were assessed using the Qubit RNA Assay Kit (Life Technologies Corp., Carlsbad, CA) and the Agilent 2100 Bioanalyzer with the RNA Nano kit (Agilent Technologies, Santa Clara, CA). Only samples with an RNA Integrity Number (RIN) ≥ 8 were used. Libraries were constructed using the TruSeq RNA Sample Prep Kit v2 according to the manufacturer’s reference guide (Illumina, Inc., San Diego, CA). Briefly, poly-A-containing RNA was purified from 500 ng of total RNA using oligo-dT-attached magnetic beads, then fragmented in a divalent cation solution at 94°C for 4 minutes. The fragmented RNA was converted into cDNA using reverse transcriptase and random hexamers, followed by second-strand cDNA synthesis. The double-stranded cDNA was end-repaired, adenylated at the 3’ ends, and ligated with indexing adapters. The prepared libraries were then purified and enriched by PCR to create the final library. The quantity and average fragment length of the final libraries were assessed using the Agilent Bioanalyzer DNA 7500 Assay (Agilent Technologies) and the Qubit High Sensitivity DNA Assay (Life Technologies Corp.). All libraries were normalized to 10 nM in Buffer EB (Qiagen, Santa Clarita, CA) and pooled in equal molar amounts. Sequencing was performed on two lanes of an Illumina HiSeq 2500 in high output mode using 100-bp paired-end reads, as per the manufacturer’s instructions. An average of 32 million paired reads were obtained per replicate. The RNAseq data has been deposited into the Sequence Read Archive with the BioProject accession code PRJNA1201272.

### 2.4 Data analysis

RNAseq data were aligned to the reference genome Hg38 using the STAR aligner. The counts obtained were normalized using DESeq2 and processed for differential analysis.

Principal Component Analysis (PCA) was performed on the variance-stabilized expression matrix to summarize the major axes of transcriptional variation across samples. PCA reduces high-dimensional gene expression data into orthogonal components that capture maximal variance, enabling visualization of global transcriptome structure. The first two principal components (PC1 and PC2) were visualized using a PCA biplot generated with the FactoMineR and factoextra R packages, with gene loading vectors displayed to highlight the features most strongly contributing to separation along PC1 and PC2. Samples were colored by genotype and treatment group to assess biological clustering and identify potential batch effects.

To further investigate nonlinear sample relationships and local neighborhood structure within the transcriptome, we applied t-Distributed Stochastic Neighbor Embedding (t-SNE) using the top 20 principal components as input. t-SNE was computed using the Rtsne package, and visualized with ggplot2. A perplexity value of 6 was selected based on recommended ranges proportional to sample size, balancing local and global structural preservation.

Gene set enrichment analysis was performed using the GOAT [40] R package (v1.0) within a set of custom in-house R scripts developed to automate enrichment testing, visualization, and comparative analysis across experimental conditions. GOAT (Gene set Ordinal Association Test) is a preranked gene set enrichment algorithm that utilizes squared rank-transformed effect sizes to compute gene scores and evaluates significance against precomputed skew-normal null distributions, enabling parameter-free and computationally efficient testing.

Differential expression outputs (log₂ fold changes) were supplied to GOAT as pre-ranked gene lists. Gene Ontology annotations were derived from the January 1, 2025 Gene Ontology release (*go_2025-01-01.obo.gz*), and enrichment analyses were performed separately across the Biological Process (GO_BP), Molecular Function (GO_MF), and Cellular Component (GO_CC) branches. Multiple-testing correction was applied using Bonferroni adjustment within each ontology domain and further corrected for testing across three GO domains. In addition, Oncogenic Signature and Hallmark pathway enrichments were evaluated using gene sets from the MSigDB GSEA collection v2025.1, processed through the same GOAT-based pre-ranked framework.

#### 2.4.1 Machine-Learning Identification of Discriminant Gene Signatures

To identify genes most strongly associated with treatment-dependent transcriptional divergence, we applied two complementary supervised classification approaches—Random Forest (RF)[41] and Partial Least Squares-Discriminant Analysis [42] (PLS-DA)—using log1p-transformed normalized expression values from MDA-MB-157 and APC-knockdown cell lines exposed to cisplatin, paclitaxel, or untreated control conditions.

Random Forest models were constructed using 2,000 trees with balanced bootstrapping of samples and randomized feature selection at each split. Feature importance was evaluated by Mean Decrease in Accuracy, representing reduction in predictive performance when each gene is permuted. The top-ranked genes were selected based on highest importance values.

PLS-DA was performed using in-house R scripts using the mixOmics package [43] with two latent components optimized via classification error minimization. Gene contribution was quantified using VIP (Variable Importance in Projection) scores, with the top features representing those genes most responsible for supervised group separation. To derive a stable and biologically interpretable discriminant signature, we extracted the intersection of the top RF and PLS-DA gene sets, reducing methodological bias and increasing reliability. Candidate signature genes were visualized by hierarchical clustering heatmaps (z-scored rows), and functional relevance was evaluated through literature and pathway enrichment.

## 3. Results and Discussion

MDA-MB-157 cells were originally derived from the pleural effusion of an African American patient with metastatic breast cancer. They are classified as triple-negative breast cancer (TNBC) cells, meaning they lack the expression of estrogen receptors (ER), progesterone receptors (PR) and HER2 receptors, making them a model for studying aggressive and difficult to treat forms of breast cancer. We used RNA-seq to profile the genome-wide expression of genes in MDA-MB-157 and APC knockdown cells with and without drug treatment (PTX and CIS).

Principal Component Analysis (PCA) and t-Distributed Stochastic Neighbor Embedding (t-SNE) were used to assess global transcriptional relationships among APC_shRNA1, APC_shRNA2, and parental MDA-MB-157 samples under control (CON), cisplatin (CIS), and paclitaxel (PTX) conditions (Figure 1). These dimensionality-reduction analyses showed that PCA provided only partial separation of groups, whereas t-SNE achieved more distinct clustering by genotype and chemotherapeutic treatment, highlighting nonlinear transcriptomic variation among samples. Together, these analyses confirm complex genotype-driven transcriptomic divergence and treatment-associated transcriptional shifts, providing a foundation for downstream differential expression and pathway enrichment analyses.

**Figure 1.**
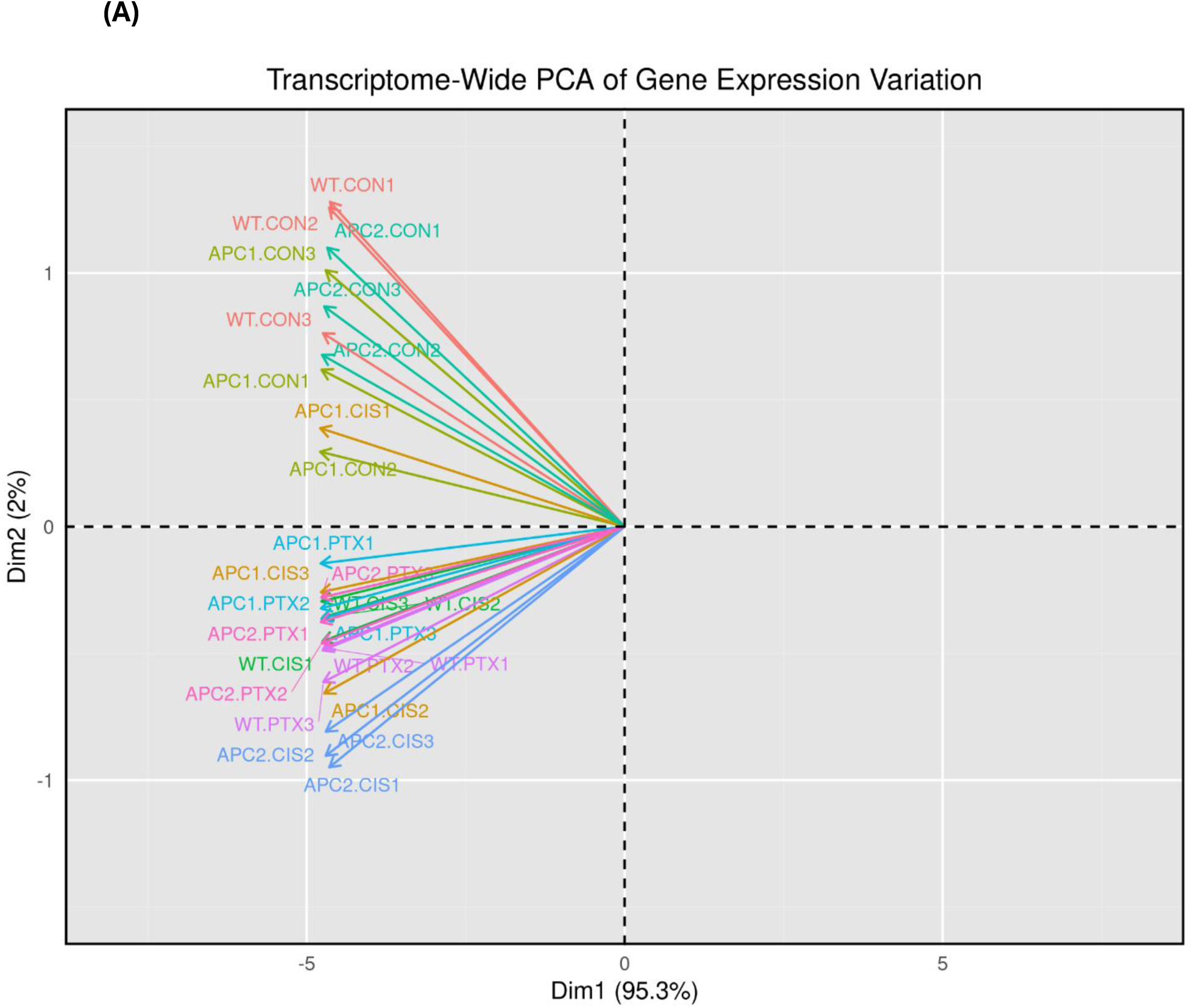
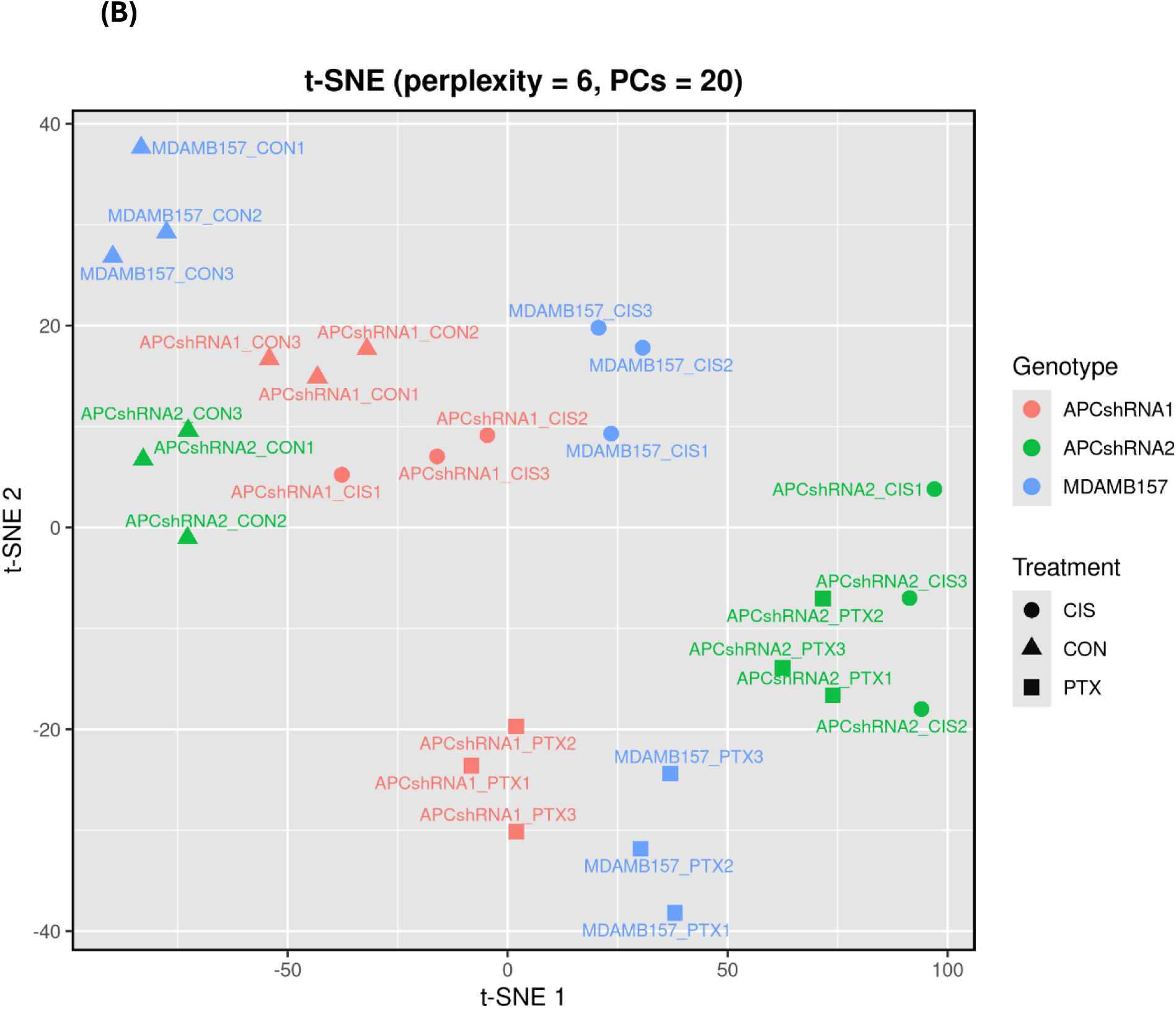
Global transcriptome structure visualized by PCA and t-SNE. **(A) Principal Component Analysis (PCA) biplot** of global gene-expression profiles from APC knockdown (APC_shRNA1, APC_shRNA2) and parental MDA-MB-157 cells under control (CON), cisplatin (CIS), and paclitaxel (PTX) treatment conditions. PC1 explains **95.3%** of total variance and separates samples primarily by genotype, while PC2 explains **2%** and contributes to treatment-based separation. Vectors represent gene loading directions, indicating the transcriptional features that drive major axes of variation. **(B) t-Distributed Stochastic Neighbor Embedding (t-SNE) plot** showing nonlinear separation of the same transcriptomic profiles using perplexity = 6 and the top 20 principal components as input. Samples cluster distinctly by genotype (APC_shRNA1, APC_shRNA2, MDA-MB-157) and separate further by treatment status (CON, CIS, PTX).

The transcriptomic data was then analyzed using GOAT [40]. The data was not filtered based on log fold change. This approach not only considers genes that are significantly changed but also includes smaller changes in gene expression of multiple genes within the context of a pathway or biological process. This method acknowledges that a single significantly dysregulated gene in a pathway may not impact the pathway as much as several dysregulated genes, which may provide more flux to the pathway or the affected signaling system. Further, GOAT analysis was done for enrichment of oncogenic signature gene sets as they represent the transcriptional responses to activation or inhibition of oncogenes and tumor suppressors. They are derived from experiments where specific genes known to be involved in cancer are manipulated, such as by overexpression or knockdown, and the resultant changes in gene expression are analyzed.

In the following paragraphs, we will first systematically describe the results of enriched signatures transcriptome, followed by the analysis of the discriminant signatures obtained using machine learning analysis.

### 3.1 Baseline transcriptional rewiring in APC knockdown lines relative to parental MDA-MB-157 cells

To determine the impact of APC loss on global pathways prior to chemotherapeutic exposure, we compared transcriptomic profiles of APCshRNA1 and APCshRNA2 to parental MDA-MB-157 cells under baseline (no-drug) conditions. Gene set enrichment analysis integrating GO (BP/MF/CC), Hallmark pathways, and Oncogenic Signatures revealed broad remodeling of cellular identity, stress-response programs, and metabolic architecture in both knockdown lines; however, each displayed a distinct enrichment landscape consistent with its experimentally confirmed drug-resistance phenotype [38].

#### 3.1.1 Comparison of Enriched signatures between MDA-MB-157 and APC Knockdown Cells

APC_shRNA1 exhibited marked enrichment of Hallmark pathways related to epithelial–mesenchymal transition (EMT), inflammatory NF-κB–driven survival signaling, enhanced mitochondrial oxidative phosphorylation, cellular stress responses, and interferon-alpha signaling, indicating a transcriptional state consistent with increased motility, stress tolerance, and metabolic flexibility.

GO Biological Process (GO-BP) enrichment indicated upregulation of pathways involved in regulation of transcription and RNA metabolism, translation/cytoplasmic translation, and apoptotic process, together with cilium organization and signal-transduction regulation, reflecting cytoskeletal remodeling typical of EMT [44, 45]. GO Molecular Function (GO-MF) terms including chromatin binding, transcription regulator activity, cadherin binding, and structural constituent of ribosome supported broad transcriptional and translational reprogramming. GO Cellular Component (GO-CC) demonstrated strong enrichment for mitochondrial inner membrane, ribosomal subunits, ribonucleoprotein complexes, focal adhesion, anchoring junction, and external plasma membrane—consistent with OXPHOS upregulation and adhesion remodeling [46, 47]. Oncogenic signaling patterns indicated a transcriptional program marked by proliferative competence, epithelial–mesenchymal transition (EMT), growth-factor–driven inflammatory survival signaling, and MAPK/ERK pathway modulation. Together with evidence of increased mitochondrial oxidative metabolism, chromatin remodeling, and enhanced focal-adhesion and cytoskeletal dynamics, these features define a metabolically adaptable, motile, and anti-apoptotic phenotype. This integrated signature is consistent with the experimentally observed paclitaxel-tolerant behavior of APC_shRNA1 cells [38] (see Figure 2(A), and Supp1_APCshRNA1_CON_vs_MDAMB_CON.xlsx).

**Figure 2(A).**
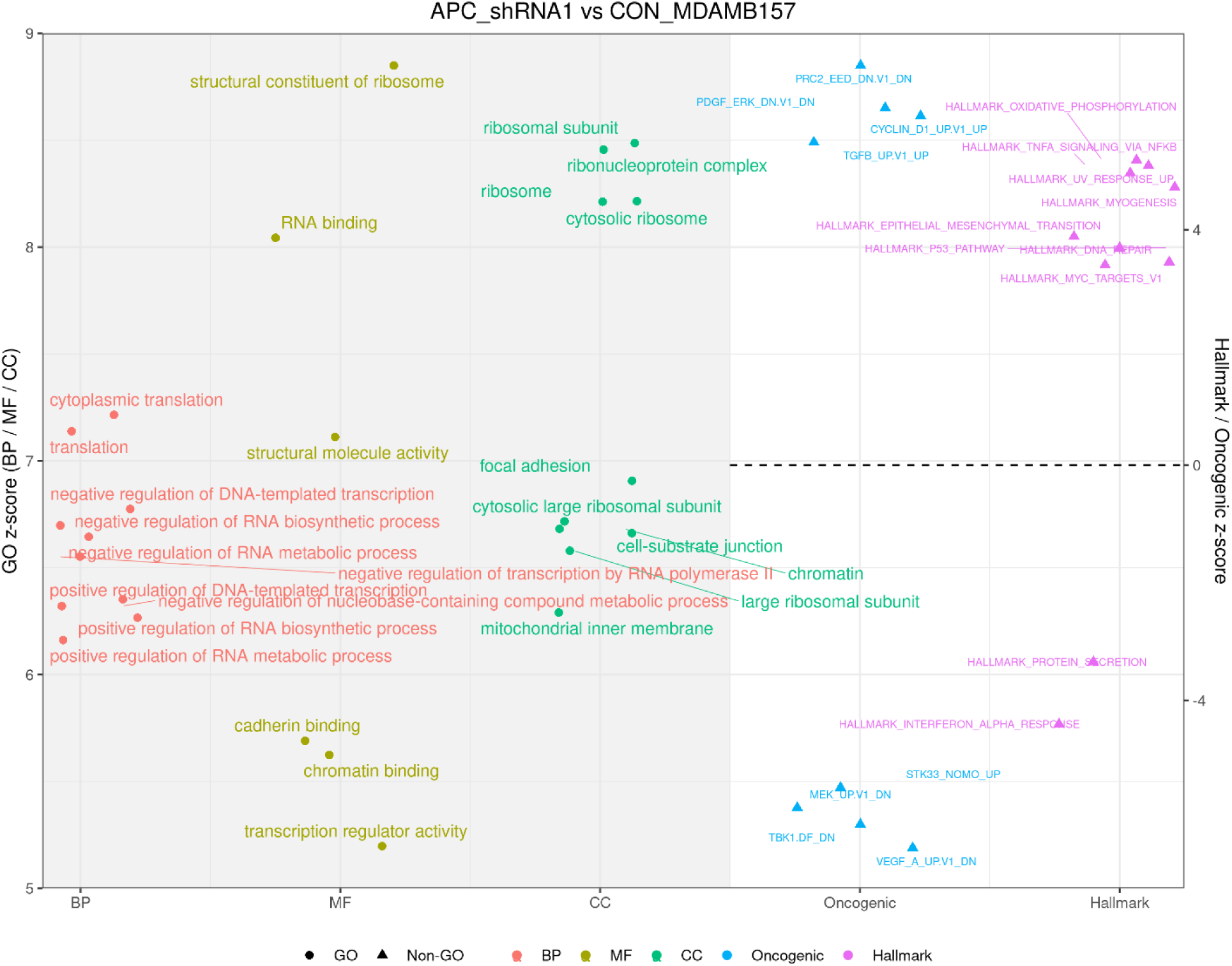
APC_shRNA1 vs CON_MDAMB157: Scatter plot showing the top enriched GO, Hallmark, and Oncogenic pathways distinguishing APC_shRNA1 from parental MDA-MB-157 control cells. APC_shRNA1 displays prominent enrichment of RNA biosynthesis and translational machinery, ribosomal structure and function, mitochondrial inner-membrane metabolism, focal adhesion and cytoskeletal remodeling, and chromatin regulatory programs. Hallmark and Oncogenic categories support activation of inflammatory NF-κB/TGF-β signaling, EMT remodeling, oxidative phosphorylation, and cell-cycle–associated pathways, reflecting a metabolically active, proliferative, and motile phenotype aligned with survival under cytotoxic stress.

APC_shRNA2 exhibited a distinct baseline enrichment pattern characterized by enhanced antioxidant and redox-buffering capacity, preparedness for DNA damage response, and extracellular matrix and structural remodeling. The pathway profile included strong activation of inflammatory and immune response programs, including TNF-α/NF-κB signaling and type I and II interferon responses, together with heme and iron metabolic regulation, increased mitochondrial oxidative phosphorylation, and epithelial–mesenchymal transition (EMT)-associated transcriptional remodeling [48, 49]. These transcriptional features imply that APC_shRNA2 cells adopt a pre-conditioned stress-resilient state with increased capacity to withstand genotoxic and cytotoxic stimuli, providing a mechanistic basis for their potential to better tolerate chemotherapy-induced damage relative to parental cells (see Figure 2(B) and Supp2_APCshRNA2_CON_vs_MDAMB_CON.xlsx).

**Figure 2(B).**
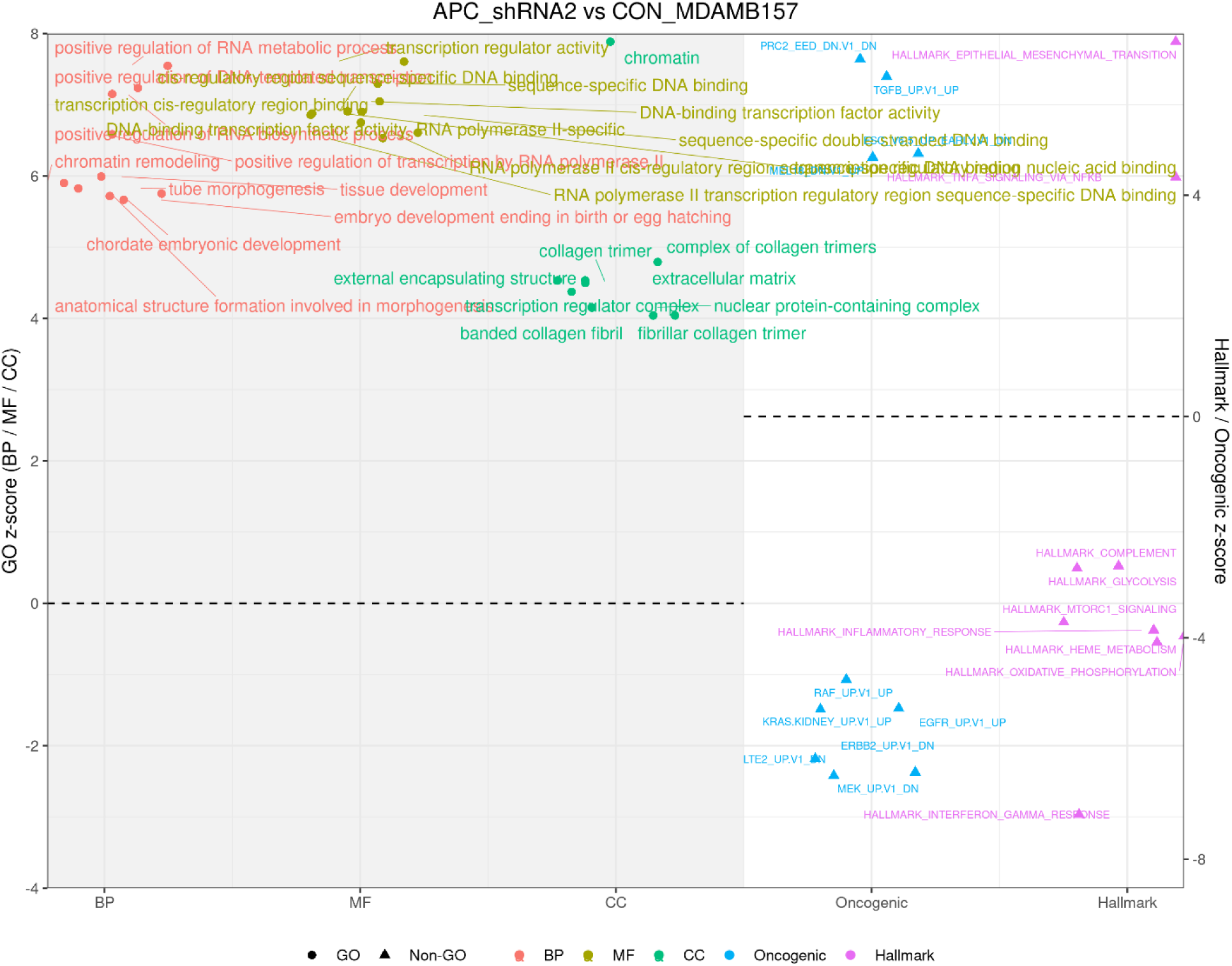
APC_shRNA2 vs CON_MDAMB157: Scatter plot summarizing the top enriched pathways from Gene Ontology (GO: BP, MF, CC) and Hallmark/Oncogenic signature categories comparing APC_shRNA2 to parental MDA-MB-157 control cells. Positive z-scores indicate pathway enrichment relative to controls. GO pathways (left, grey-shaded region) reveal significant upregulation of transcription factor regulation, chromatin remodeling, RNA metabolism, extracellular matrix remodeling, and collagen organization. Hallmark and Oncogenic pathways (right, white region) show strong activation of inflammatory signaling, interferon responses, metabolic and oxidative phosphorylation programs, and epithelial–mesenchymal transition (EMT), consistent with elevated stress resilience and transcriptional plasticity.

GO-BP analysis highlighted enrichment for developmental processes, chromatin organization, and positive/negative regulation of transcription and RNA biosynthesis, consistent with transcriptional plasticity. GO-MF revealed increased DNA-binding transcription factor activity, chromatin binding, double-stranded DNA binding, and actin-filament binding, while GO-CC demonstrated enrichment of chromatin complexes and collagen-trimer / ECM structures, reflecting both nuclear epigenetic control and extracellular adhesion remodeling.

Oncogenic signaling analysis revealed enrichment of pathways associated with EGFR- and KRAS-mediated growth signaling, RAF/MAPK cascade activation, TGF-β– driven EMT and remodeling, and NRF2-regulated antioxidant defense, along with reduced PRC2/EED regulatory activity affecting chromatin accessibility. Together, these features support activation of pro-survival and inflammatory signaling networks, enhanced redox buffering capacity, and epigenetic plasticity [50, 51]. This profile reflects a transcriptional state that is redox-protected and primed for DNA repair, with reinforced cytokine/NF-κB signaling, extracellular matrix and adhesion remodeling, and chromatin-mediated adaptability—hallmarks commonly associated with cisplatin tolerance [52].

### 3.2 Profiling Enrichment in MDA-MB 157 Under Chemotherapeutic Stress

Cisplatin and paclitaxel are both commonly used chemotherapeutic agents in the treatment of triple-negative breast cancer (TNBC). Considering that MDA-MB-157 is a TNBC cell line, understanding the effects of these drugs on this model is crucial.

#### 3.2.1 Comparison of Enriched Hallmark Gene Sets and GO Terms in MDA-MB-157 Under Cisplatin Treatment

Gene set enrichment analysis revealed that cisplatin exposure induced a coordinated transcriptomic response characterized by strong activation of Hallmark pathways associated with DNA damage repair and cell-cycle checkpoint programs (Figure 3(A) and Supp3_MDAMB_CIS_vs_MDAMB_CON.xlsx). Pathways associated with genome integrity—including DNA repair mechanisms and p53-mediated checkpoint control—were prominently enriched, accompanied by signatures indicating reinforced chromosome segregation, microtubule stability, and mitotic spindle organization. Rather than undergoing complete proliferative collapse, cells appeared to fortify spindle architecture and maintain checkpoint-governed cycling, consistent with a damage-tolerant proliferative state.

**Figure 3(A).**
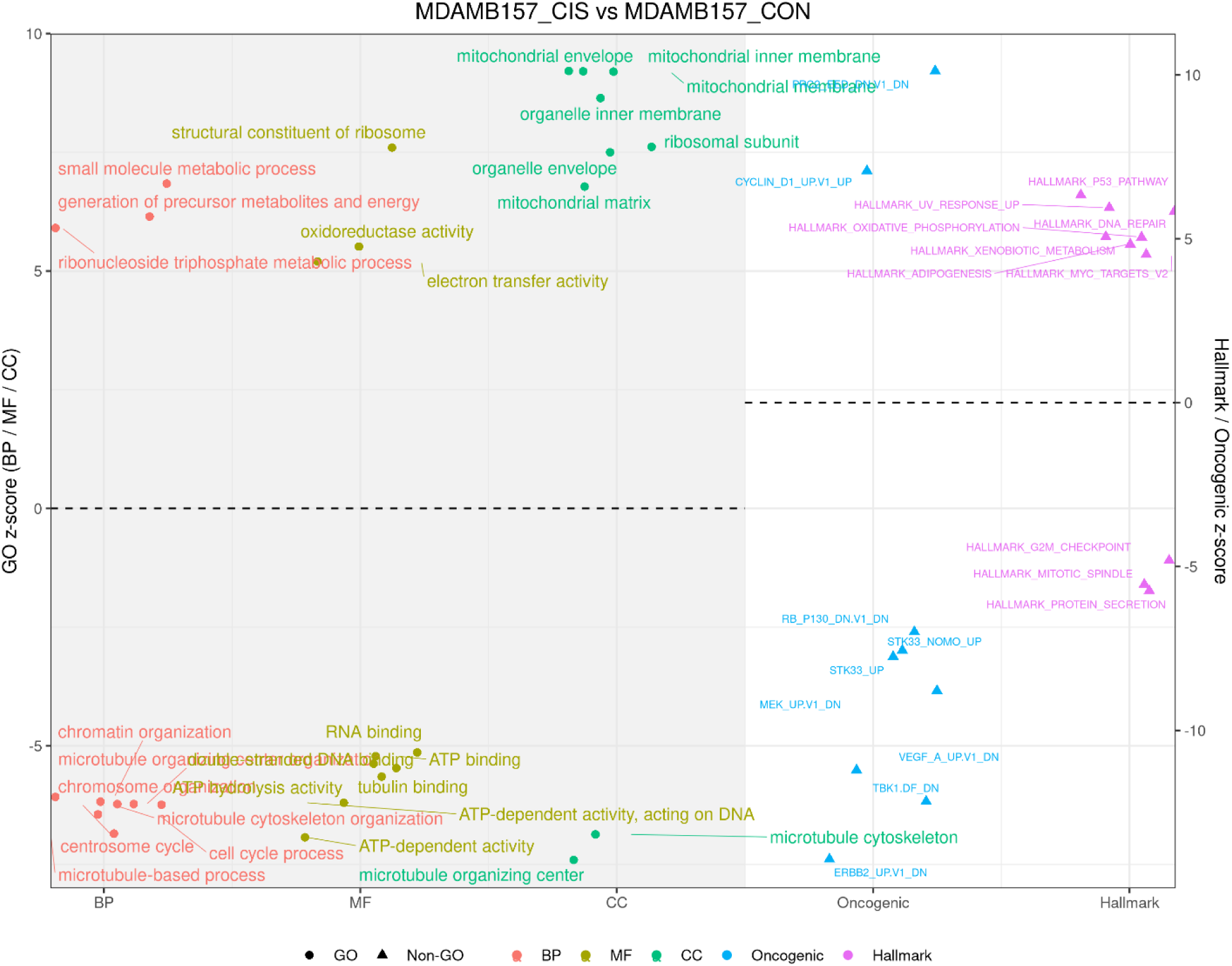
MDAMB157_CIS vs MDAMB157_CON: Scatter plot summarizing the top enriched Gene Ontology (GO) Biological Process (BP), Molecular Function (MF), and Cellular Component (CC) categories, along with Hallmark and Oncogenic signature pathways, comparing cisplatin-treated MDA-MB-157 cells to untreated controls. GO terms (left panels, grey background) show strong positive enrichment of mitochondrial organization, respiratory chain activity, electron transfer processes, and ribosomal structure, indicating mitochondrial bioenergetic and translational reinforcement under DNA-damaging stress. Among Hallmark pathways (right panel), DNA damage repair, oxidative phosphorylation, p53 checkpoint signaling, and UV stress response modules are significantly enriched, consistent with canonical cisplatin-induced DNA damage responses. Several mitotic-associated pathways, including G2/M checkpoint, mitotic spindle, and protein secretion, are downregulated (negative z-scores), reflecting checkpoint arrest and suppression of mitotic progression. Blue oncogenic signature markers display negative z-scores, indicating reduced MAPK-driven proliferative signaling under cisplatin.

Metabolic and organellar remodeling also featured prominently. GO Enrichment of mitochondrial oxidative phosphorylation, mitochondrial inner-membrane and matrix components, and nucleotide/ATP biosynthetic processes indicated a shift toward high-efficiency energy production to support the substantial metabolic demands of DNA lesion processing and recovery. Concurrent up-regulation of translational and proteostasis pathways—including ribosomal assembly, translation initiation, protein secretion, and RNA metabolism—suggested increased protein synthesis capacity to support stress response execution.

Cytoskeletal and structural remodeling pathways—including microtubule and tubulin-binding functions, spindle and kinetochore complexes, and focal adhesion components—were strongly represented, indicating stabilization of the division machinery during chemotherapeutic stress. Interferon and innate inflammatory signaling modules were also activated, consistent with stress-induced immune-like signaling and possible cGAS–STING pathway engagement.

Finally, oncogenic program dynamics revealed reinforcement of proliferative drivers such as E2F, MYC, and Cyclin-D–linked transcriptional activity, while signaling associated with receptor-tyrosine-kinase/MAPK pathways (including EGFR/MEK/RAF axes) appeared relatively reduced. This suggests a shift away from growth-factor dependency and toward a DNA-damage-response–centered survival strategy.

Collectively, these findings support a mechanistic model in which cisplatin induces a repair-committed, biosynthetically and metabolically engaged state that preserves proliferative capacity under checkpoint control, stabilizes cytoskeletal machinery, enhances proteostasis, and recruits innate stress signaling. Such coupling of DDR, OXPHOS-based metabolic support, translation, and spindle reinforcement represents a clinically relevant path toward damage-adaptive persistence and residual disease following genotoxic therapy[46, 47].

#### 3.2.2 Comparison of Enriched Hallmark Gene Sets and GO Terms in MDA-MB-157 Under Paclitaxel Treatment

Gene set enrichment analysis revealed that paclitaxel treatment induces a coordinated adaptive response across DNA damage signaling, metabolic support systems, translational capacity, inflammatory survival signaling, and chromatin remodeling, while simultaneously repressing pathways associated with mitotic spindle assembly. Notably, the mitotic spindle program was down regulated, reflecting transcriptional suppression of spindle-structural genes. This downregulation is consistent with paclitaxel’s mechanism of action, whereby microtubule stabilization disrupts spindle dynamics, activates the spindle assembly checkpoint, and prevents proper chromosomal segregation (Figure 3B and Supp4_MDAMB_PTX_vs_MDAMB_CON.xlsx).

**Figure 3(B).**
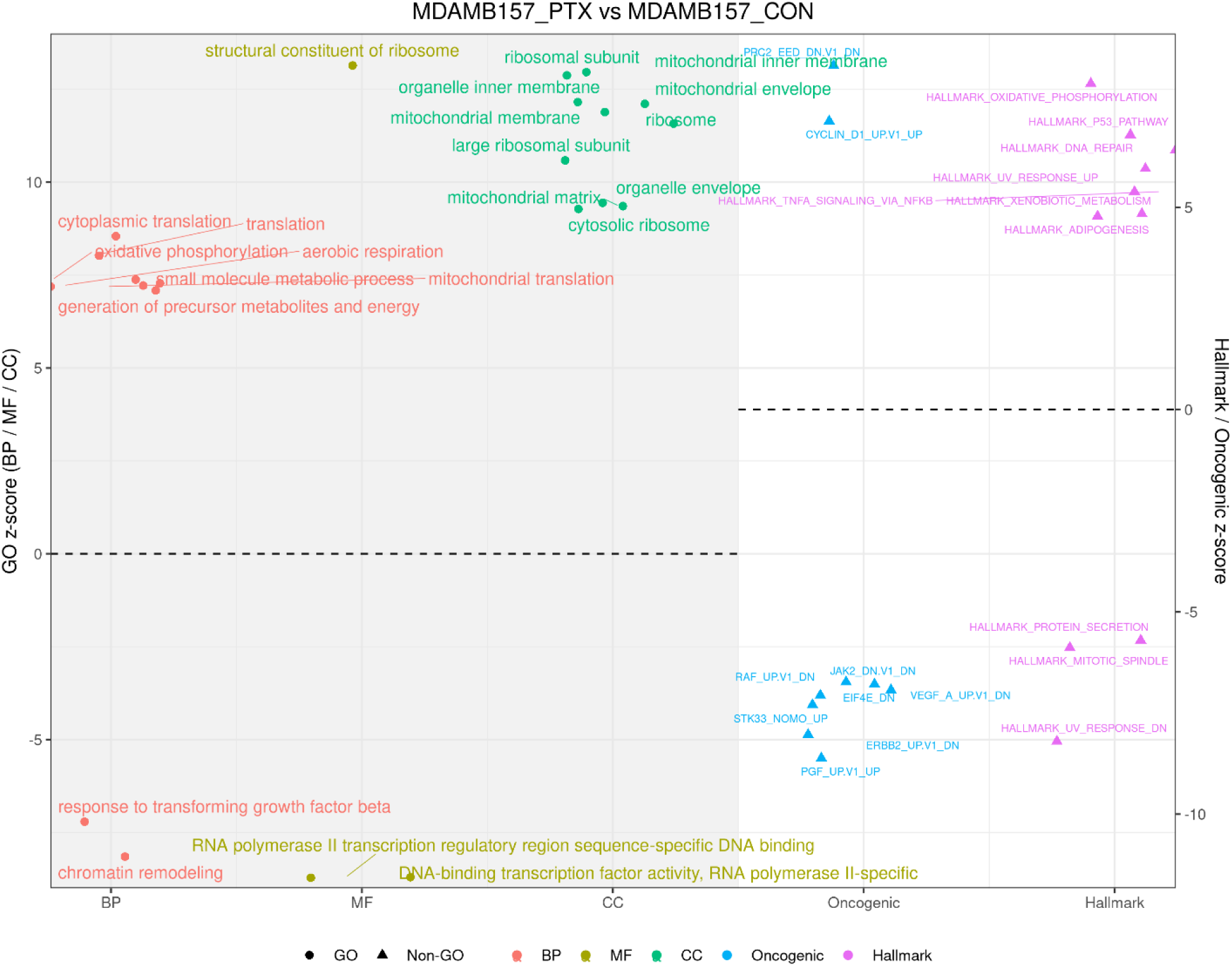
MDAMB157_PTX vs MDAMB157_CON: Scatter plot showing pathway enrichment following paclitaxel treatment. Similar to cisplatin, GO terms indicate strong enrichment of mitochondrial metabolic pathways, ribosomal structure, electron transport, and ATP-producing metabolism, demonstrating substantial bioenergetic engagement. However, Hallmark analysis reveals notable downregulation of mitotic spindle and protein secretion pathways (large negative z-scores), consistent with microtubule stabilization and spindle checkpoint activation induced by paclitaxel. Inflammatory TNF-α/NF-κB signaling, oxidative phosphorylation, DNA repair, and MYC-linked proliferative programs are positively enriched, indicating a biosynthetically active survival response. Oncogenic signaling shows broad downregulation of MEK/RAF/ERBB2 pathways, suggesting a shift away from canonical MAPK dependence. Overall, paclitaxel induces checkpoint activation, metabolic reinforcement, and adaptive translational and inflammatory responses supporting treatment tolerance.

Despite spindle pathway repression, paclitaxel-treated cells demonstrated strong enrichment of p53-mediated checkpoint signaling, DNA repair programs, oxidative phosphorylation–linked mitochondrial metabolism, protein secretion, and reactive oxygen species detoxification pathways, indicating a biosynthetically active, energy-intensive survival state. GO Biological Process categories highlighted mitochondrial electron transport and translation, ATP and nucleotide biosynthesis, RNA metabolism, and TGF-β pathway signaling, supporting a stress-adapted, repair-competent phenotype. Corresponding GO Cellular Component enrichments included respiratory chain complexes, mitochondrial and cytosolic ribosomes, and microtubule apparatus regulators, while molecular function categories revealed increases in histone kinase activity, electron transfer capacity, and tubulin-binding functions.

Inflammatory signaling modules such as TNF-α/NF-κB and oxidative detoxification programs were strongly activated, consistent with pro-survival inflammatory circuitry that buffers apoptosis. Oncogenic signaling dynamics indicated a shift away from MEK/RAF/ERBB2 dependency toward EGFR/SRC/MYC/EIF4E-linked translational control, accompanied by chromatin derepression signatures suggestive of enhanced transcriptional plasticity [44].

### 3.3 Profiling Enrichment in APC sh-RNA1 and APCshRNA2 Under Chemotherapeutic Stress

Since APC plays a significant role in TNBC by regulating key cellular processes essential for cancer progression and response to therapy, and considering that APC is frequently lost in TNBC cases, two APC knockdown models of MDA-MB157, APC sh-RNA1 and APCshRNA2 were studied for transcriptomic changes in response to paclitaxel and cisplatin exposure.

#### 3.3.1 Comparison of Enriched Hallmark Pathways, GO Ontologies, and Oncogenic Signatures in APC-shRNA1 Under Cisplatin Treatment

Pathway enrichment analysis revealed that cisplatin exposure in APC_shRNA1 cells induces a coordinated transcriptional program centered on DNA damage surveillance, metabolic reinforcement, and translational capacity. Hallmark pathways demonstrated strong activation of p53 signaling, DNA repair machinery, G2/M checkpoint control, and mitotic spindle-associated processes, indicating robust engagement of genome integrity and checkpoint systems under genotoxic stress. Concurrent enrichment of oxidative phosphorylation, fatty-acid and xenobiotic metabolism, and reactive oxygen species detoxification suggests a shift toward mitochondrial ATP production and redox buffering to meet the high energetic and oxidative demands of DNA repair. Elevation of MYC-associated transcriptional programs and stress-response modules, including interferon and UV-induced signaling, reflects broad transcriptional reprogramming and innate-like stress adaptation (Figure 4(A) and Supp5_APCshRNA1_CIS_vs_APCshRNA1_CON.xlsx).

**Figure 4(A).**
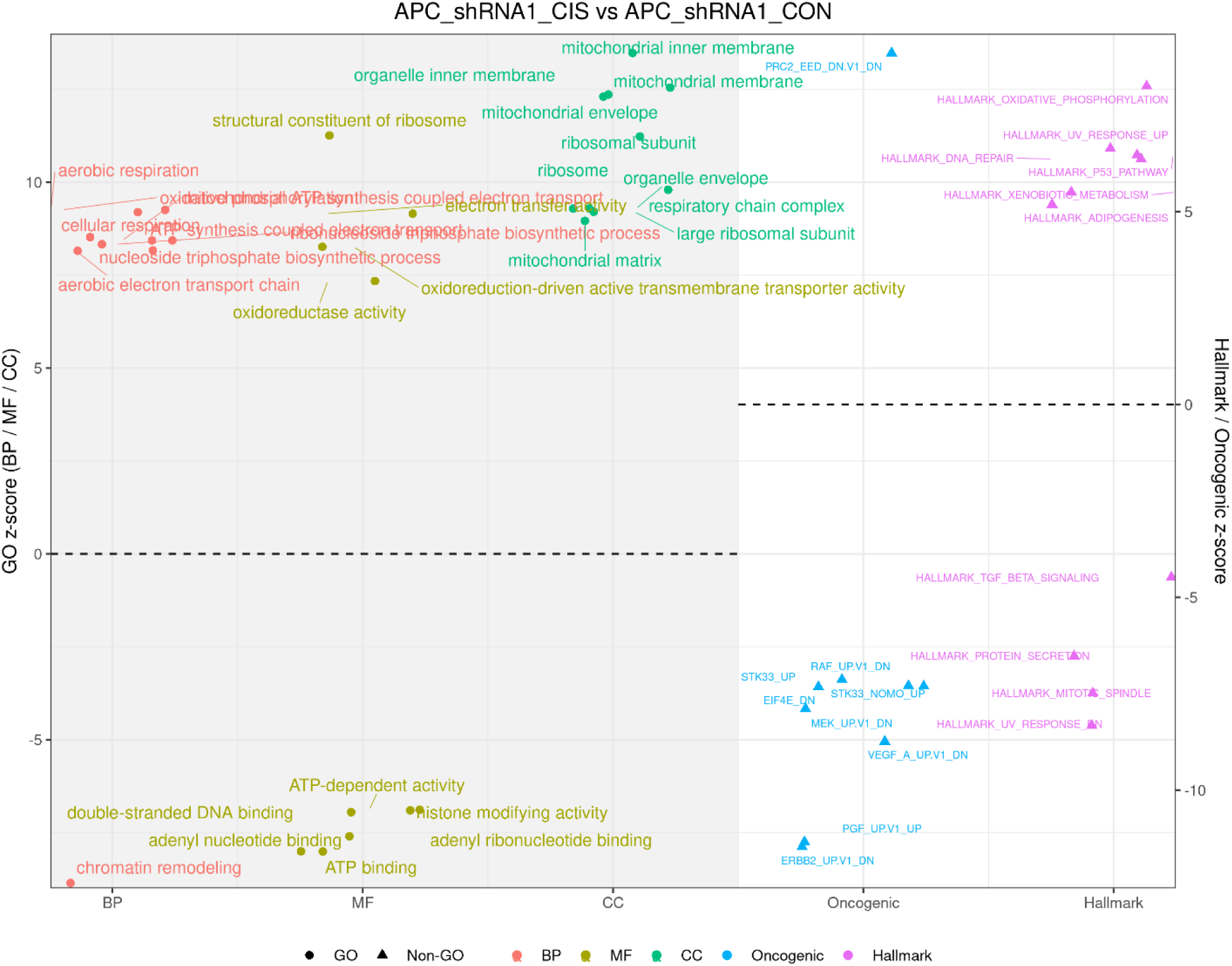
Enrichment of Hallmark, Oncogenic Signatures, and GO pathways in APC_shRNA1 cells treated with cisplatin compared to APC_shRNA1 controls: Scatter plot showing pathway-level z-score enrichment, with GO terms (BP, MF, CC) displayed as circles and Hallmark/Oncogenic signatures as triangles. Positive z-scores indicate upregulation and negative scores indicate suppression under cisplatin treatment. Upregulated pathways include P53 signaling, DNA repair, oxidative phosphorylation, and mitochondrial and ribosomal structural components, reflecting a DNA-damage–responsive, metabolically reinforced state. Downregulated modules include Mitotic Spindle, TGF-β signaling, and RTK/MAPK oncogenic programs, consistent with checkpoint-mediated mitotic restraint and signaling rebalancing.

Complementary GO analyses supported this physiological response. Biological Process terms prominently featured oxidative phosphorylation, electron transport, ATP generation, ribonucleoside biosynthesis, and cytoplasmic translation—indicating a biosynthetically active, energy-intensive cellular state. Cellular Component categories highlighted enrichment of mitochondrial inner membrane and respiratory chain complexes, together with cytosolic and mitochondrial ribosomes and microtubule-organizing structures, consistent with enhanced metabolic output and maintenance of cytoskeletal integrity during checkpoint arrest. Molecular Function enrichment included oxidoreductase and electron-transfer activities, ATP- and histone-modifying enzyme functions, RNA-binding, and DNA-binding regulatory activity, confirming simultaneous metabolic, chromatin-remodeling, and transcriptional control processes [44, 47].

Oncogenic signaling signatures demonstrated increased activity of EGFR-, SRC-, Cyclin-D1–, EIF4E-, and MYC-driven pathways, indicating maintenance of translation and proliferative drive during treatment. Conversely, reduced MEK/RAF/ERBB2 signaling and decreased PRC2/EED-mediated repression suggest a redistribution away from RTK–MAPK dependence and toward survival circuitry supported by chromatin relaxation and transcriptional flexibility [51].

Together, these findings indicate that APC_shRNA1 cells likely respond to cisplatin by adopting an OXPHOS-powered, translation-enabled, and chromatin-flexible damage-response state, sustaining checkpoint-governed self-renewal rather than entering irreversible arrest [46]. This adaptive configuration supports tolerability to DNA-damaging stress and represents a mechanistic platform from which chemotherapeutic persistence may arise.

#### 3.3.2 Comparison of Enriched Hallmark Pathways, GO Ontologies, and Oncogenic Signatures in APC-shRNA1 Under Paclitaxel Treatment

Pathway enrichment analysis revealed that paclitaxel treatment of APC_shRNA1 cells activates a coordinated transcriptional program involving DNA damage surveillance, chromatin remodeling, mitochondrial bioenergetics, and translational capacity. Hallmark gene sets demonstrated strong enrichment of p53 signaling, DNA repair, G2/M checkpoint control, oxidative phosphorylation, ROS detoxification, xenobiotic metabolism, and protein secretion, consistent with a robust checkpoint-governed damage response accompanied by metabolic reinforcement. Notably, mitotic spindle components were significantly down-regulated, indicating checkpoint-mediated mitotic restraint rather than collapse, a characteristic adaptation to paclitaxel-induced spindle tension (Figure 4(B) and Supp6_APCshRNA1_PTX_vs_APCshRNA1_CON.xlsx).

**Figure 4(B).**
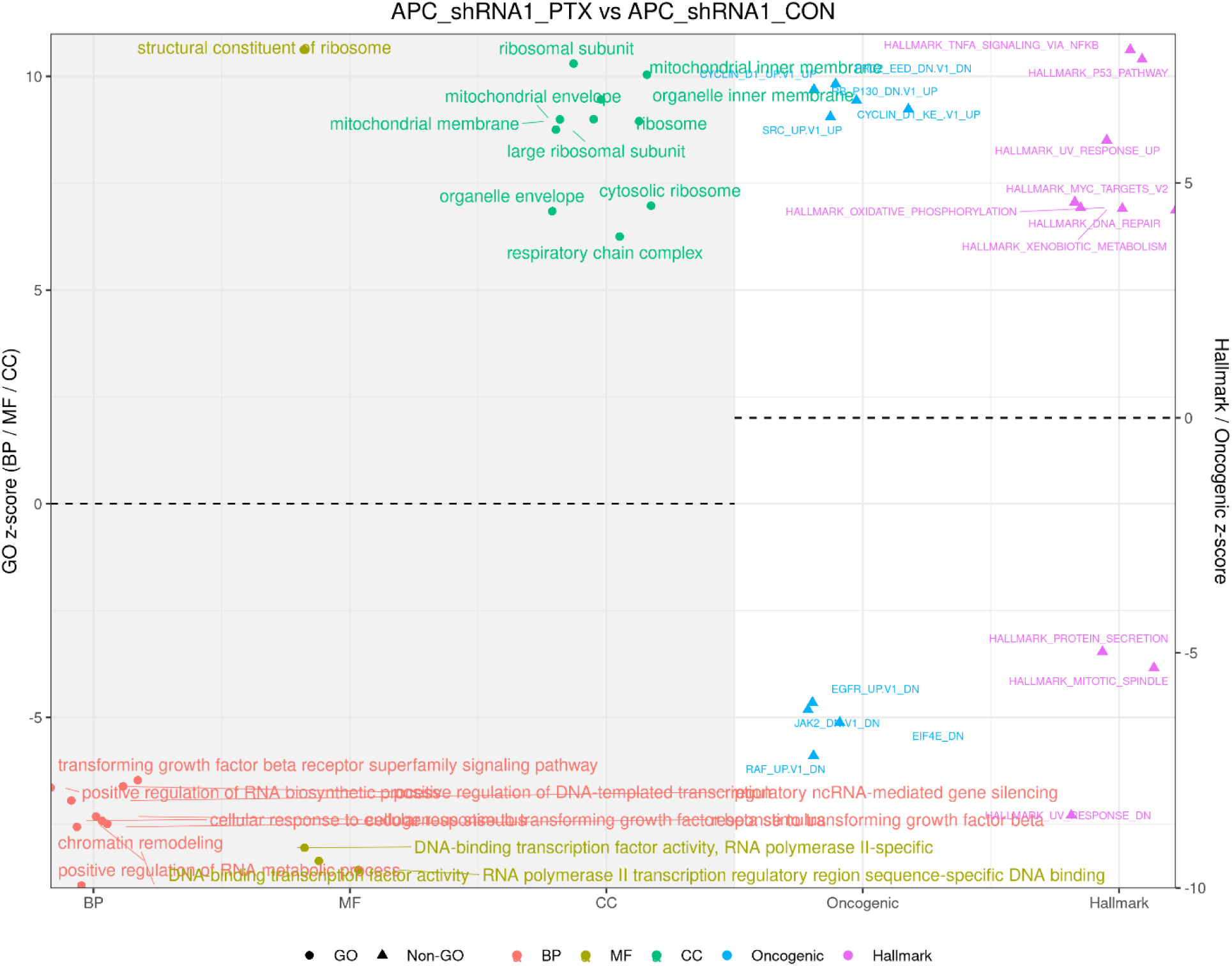
Pathway enrichment landscape in APC_shRNA1 cells treated with paclitaxel relative to untreated APC_shRNA1 controls. Scatter plot illustrating enrichment z-scores for Gene Ontology (GO) Biological Process, Molecular Function, and Cellular Component terms (circles), and Hallmark/Oncogenic Signature gene sets (triangles). Positive values indicate upregulated pathways and negative values represent downregulated pathways following paclitaxel exposure. Paclitaxel treatment strongly enhanced mitochondrial and ribosomal components, oxidative phosphorylation, p53 signaling, DNA repair, xenobiotic metabolism, and NF-κB inflammatory modules, indicating a bioenergetically supported checkpoint-regulated stress response. Suppression of Mitotic Spindle and related oncogenic MAPK components reflects paclitaxel-induced spindle checkpoint activation and signaling realignment toward translation-centric survival.

GO Biological Process enrichment highlighted chromatin organization and remodeling, DNA repair, cytoplasmic translation, electron transport chain activity, and ATP and nucleotide metabolism, defining an energy-intensive, biosynthetically active cellular state. GO Cellular Component terms revealed strong signatures from the mitochondrial inner membrane, respiratory complexes, cytosolic and mitochondrial ribosomes, and microtubule cytoskeleton, confirming engagement of both metabolic and protein-synthesis machinery. GO Molecular Function terms included elevated RNA binding, transcription factor DNA interaction, oxidoreductase activity, and ATP-dependent chromatin remodeling, consistent with expanded transcriptional control and proteostasis during stress adaptation.

Oncogenic signature enrichment demonstrated increased activity of MYC-, EIF4E-, SRC-, AKT-, VEGF-, and Cyclin-D1–linked signaling, together supporting translation-centric survival and proliferative re-entry after checkpoint arrest, while reduced MEK/RAF/ERBB2 signaling and PRC2 loss-of-repression signatures indicate a shift away from canonical MAPK dependence toward chromatin-relaxed, metabolically supported stress tolerance.

These results indicate that APC_shRNA1 cells are likely to adapt to paclitaxel-induced microtubule stress by adopting a mitochondria-powered, translation-enabled, chromatin-flexible state that facilitates checkpoint-regulated survival rather than mitotic catastrophe. This integrated metabolic, epigenetic, and translational response likely contributes to apparent paclitaxel tolerance and cellular persistence.

#### 3.3.3 Comparison of Enriched Hallmark Pathways, GO Ontologies, and Oncogenic Signatures in APC-shRNA2 Under Cisplatin Treatment

Pathway enrichment analysis revealed that cisplatin exposure in APC_shRNA2 cells induces a coordinated transcriptional program involving DNA-damage surveillance, mitochondrial bioenergetics, translational reinforcement, and chromatin remodeling. Hallmark pathway enrichment demonstrated strong activation of p53 signaling, DNA repair, G2/M checkpoint control, oxidative phosphorylation, protein secretion, and detoxification modules including ROS and xenobiotic metabolism, indicating a checkpoint-governed DNA repair response supported by enhanced mitochondrial energy production and oxidative stress buffering. Elevated MYC-regulated transcriptional programs and UV/stress response pathways further suggest transcriptional plasticity and innate-like stress signaling (Figure 5(A) and Supp7_APCshRNA2_CIS_vs_APCshRNA2_CON.xlsx).

**Figure 5(A):**
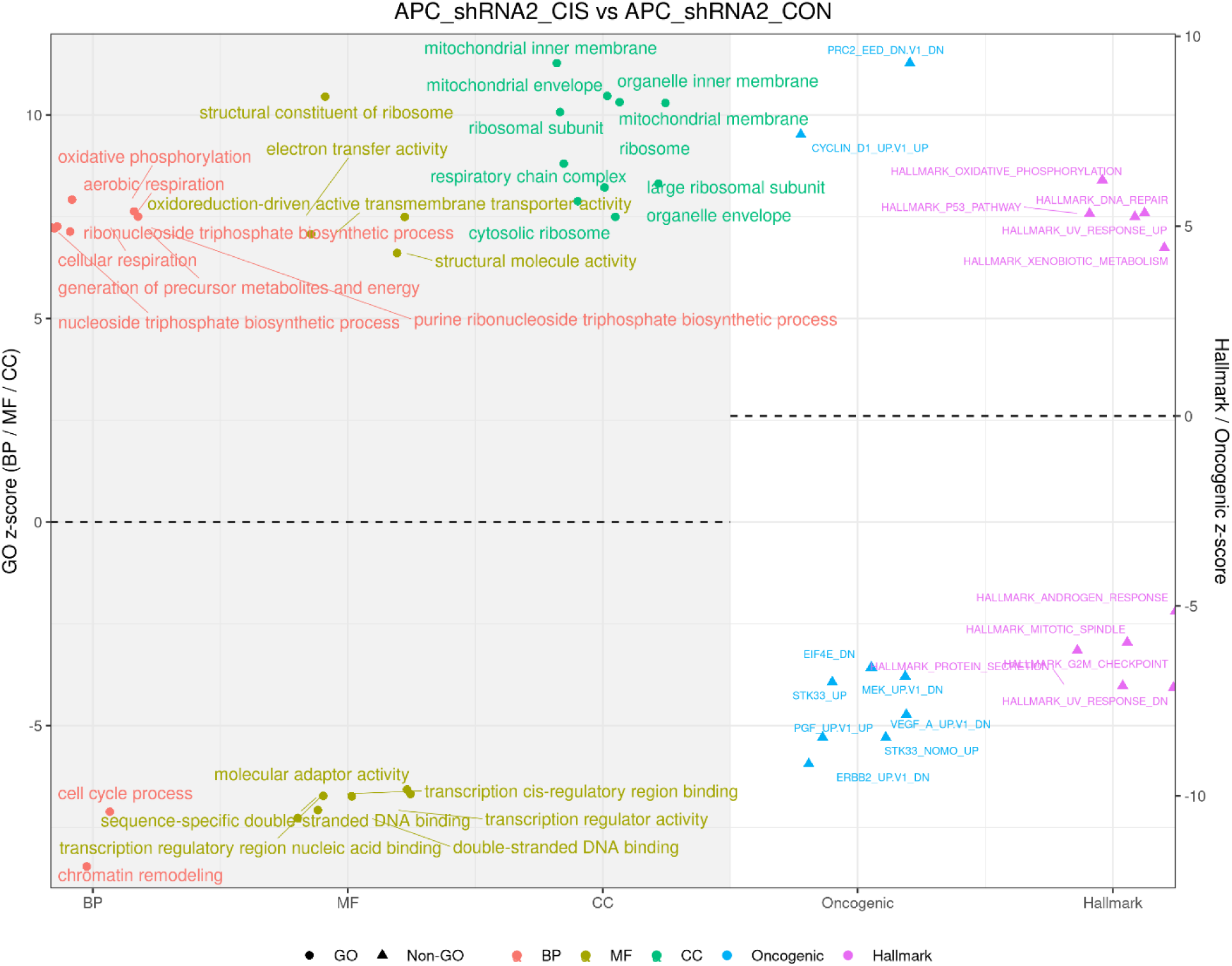
Pathway enrichment landscape in APC_shRNA2 cells treated with cisplatin relative to untreated APC_shRNA2 controls. Scatter plot illustrating z-scores for Gene Ontology (GO) Biological Process, Molecular Function, and Cellular Component terms (circles) and Hallmark/Oncogenic Signature pathways (triangles). Positive values indicate pathways upregulated following cisplatin exposure, while negative values represent downregulation. Cisplatin treatment produced strong enrichment of mitochondrial oxidative phosphorylation, respiratory chain complexes, ribosomal subunits, and nucleotide/ATP biosynthesis pathways, supporting elevated bioenergetic and translational capacity. Concurrent upregulation of p53 signaling, DNA repair, xenobiotic metabolism, and UV-response modules indicates an active DNA-damage surveillance and detoxification response. Downregulation of mitotic spindle and G2/M checkpoint signals suggests checkpoint-mediated proliferative restraint consistent with DNA-damage stress adaptation.

GO Biological Process analysis highlighted enrichment of oxidative phosphorylation, electron transport chain activity, ATP and nucleotide biosynthesis, cytoplasmic translation, and chromatin organization, identifying an energy-intensive, repair-supportive metabolic configuration. GO Cellular Component categories revealed strong signatures from the mitochondrial inner membrane, respiratory chain complexes, cytosolic and mitochondrial ribosomes, and microtubule cytoskeleton, consistent with upregulation of bioenergetic machinery, protein synthesis capacity, and reinforcement of cell-division structures under treatment. GO Molecular Function terms included electron-transfer and oxidoreductase activity, ATP binding, RNA-binding, transcription factor regulation, and histone-modifying enzyme activity, indicating extensive remodeling of metabolic, transcriptional, and chromatin regulatory circuits[47].

Oncogenic signature patterns further supported this adaptive phenotype, demonstrating increased activity of MYC-, Cyclin-D1-, EIF4E-, SRC-, and STK33-associated survival and growth effectors, while relative suppression of MEK/RAF/ERBB2 signaling and PRC2/EED-associated repression indicated a signaling shift away from canonical MAPK dependency toward translation-centric survival and epigenetically flexible transcriptional reprogramming [44].

Together, these results show that APC_shRNA2 cells adapt to cisplatin-induced damage by adopting a mitochondria-powered, translation-enabled, and chromatin-flexible stress-tolerant state, enabling sustained checkpoint-regulated survival rather than irreversible arrest. This integrated metabolic and epigenetic rewiring provides a mechanistic basis for cisplatin persistence and emergent resistance potential in APC-depleted cells.

#### 3.3.4 Comparison of Enriched Hallmark Pathways, GO Ontologies, and Oncogenic Signatures in APC-shRNA2 Under Paclitaxel Treatment

Pathway enrichment analysis revealed that paclitaxel treatment in APC_shRNA2 cells induces a coordinated adaptive program involving mitochondrial bioenergetics, translational reinforcement, chromatin remodeling, and stress-response signaling. GO Biological Process terms demonstrated strong enrichment of oxidative phosphorylation, aerobic respiration, electron transport chain function, ATP and nucleotide biosynthesis, ribosome biogenesis, post-transcriptional gene silencing, and cytoplasmic translation, defining an energy-intensive, biosynthetically active state supporting survival under spindle stress. Complementary GO Cellular Component enrichment highlighted mitochondrial inner membrane and respiratory complexes, cytosolic and mitochondrial ribosomes, and ribonucleoprotein assemblies, confirming engagement of both metabolic and translational machinery during paclitaxel response. GO Molecular Function categories further emphasized increases in RNA binding, sequence-specific DNA binding, transcriptional regulator activity, histone and chromatin-associated functions, and electron-transfer enzymatic activity, reflecting extensive restructuring of transcriptional control and chromatin accessibility (Figure 5(B) and Supp8_APCshRNA2_PTX_vs_APCshRNA2_CON.xlsx).

**Figure 5(B).**
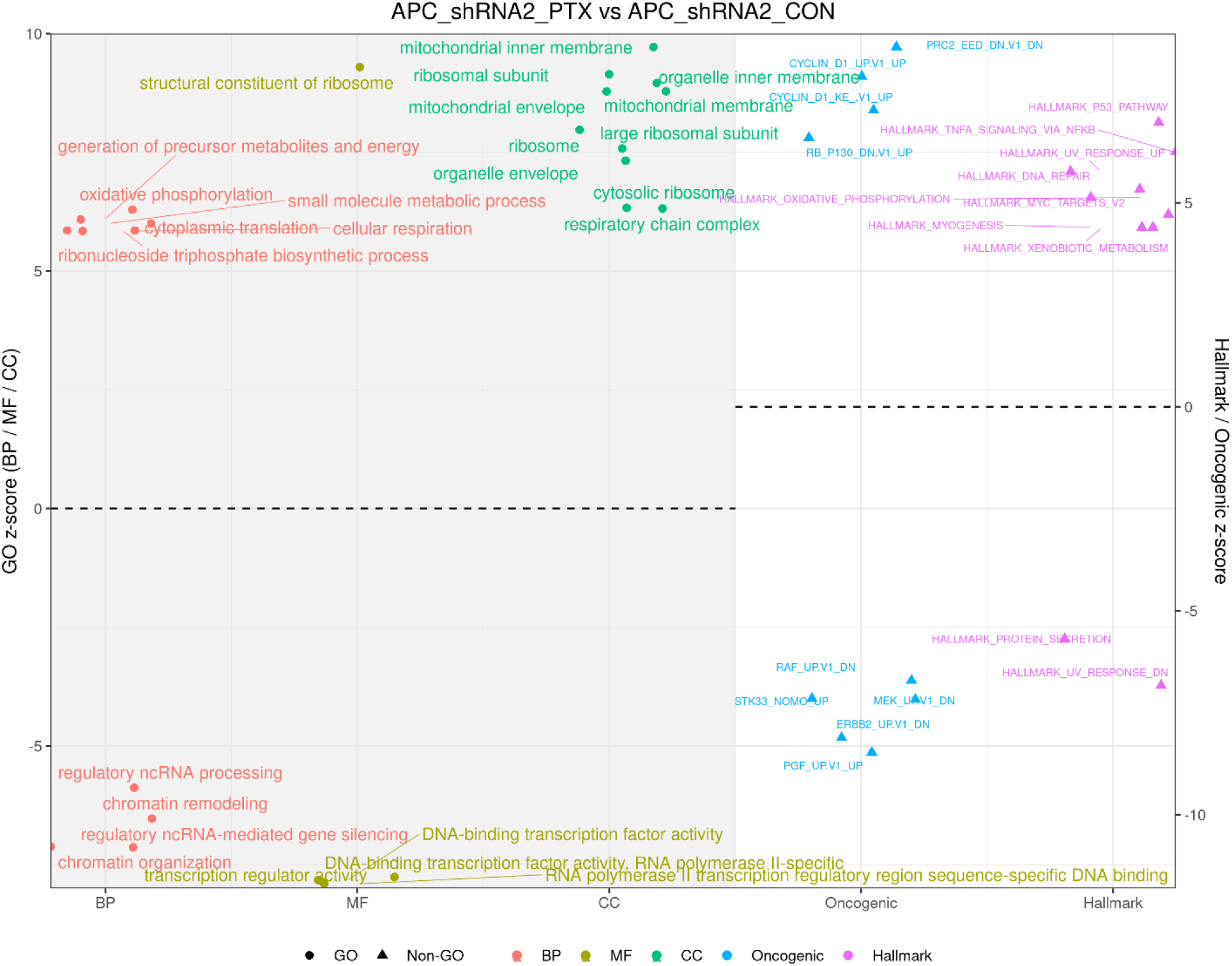
Pathway enrichment landscape in APC_shRNA2 cells treated with paclitaxel relative to untreated APC_shRNA2 controls. Scatter plot showing z-scores for enriched Gene Ontology (GO) Biological Process, Molecular Function, and Cellular Component terms (circles) and Hallmark/Oncogenic Signature pathways (triangles). Positive z-scores represent pathways upregulated after paclitaxel exposure, while negative scores represent suppression. Paclitaxel treatment strongly upregulated mitochondrial oxidative phosphorylation, respiratory chain complexes, ribosomal subunits, and nucleotide/energy metabolism pathways, indicating enhanced bioenergetic and translational capacity. Hallmark signaling modules including p53-mediated checkpoint control, DNA repair, oxidative metabolism, TNF-α/NF-κB inflammatory signaling, and xenobiotic detoxification were also elevated, reflecting an adaptive survival response. Downregulation of mitotic spindle organization and cell-cycle regulators indicates checkpoint-mediated proliferative restraint associated with paclitaxel-induced microtubule stress.

Hallmark pathway enrichment revealed strong activation of p53 signaling, DNA repair, oxidative phosphorylation, MYC-regulated programs, TNF-α/NF-κB inflammatory signaling, ROS detoxification, xenobiotic metabolism, protein secretion, and mitotic spindle stabilization, together indicating a checkpoint-regulated DNA-damage response supported by metabolic reinforcement and detoxification circuitry. Oncogenic Signature profiles showed increased activation of MYC-, AKT-, SRC-, Cyclin-D1–, EIF4E-, VEGF-associated pathways, while MEK/RAF/ERBB2 signaling was relatively reduced and PRC2/EED repression signatures appeared downregulated, suggesting a shift from classical MAPK dependency toward translation-centric survival signaling coupled to chromatin relaxation and transcriptional plasticity[44].

These results indicate that APC_shRNA2 cells adapt to paclitaxel-induced microtubule disruption by adopting a mitochondria-powered, translation-enabled, and chromatin-flexible phenotype that sustains checkpoint-governed viability and mitigates oxidative and structural stress. This adaptive rewiring supports a stress-tolerant, persistence-associated state, providing mechanistic insight into paclitaxel response behavior in APC-deficient models [23, 53].

### 3.4 Identification and Analysis of Discriminant Gene Signatures using Machine learning methods

To complement these pathway-level insights and strengthen mechanistic interpretation, we applied a supervised machine-learning feature selection strategy combining Random Forest (RF) classification and Partial Least Squares–Discriminant Analysis (PLS-DA) [41, 42]. Random Forest provides an estimate of gene importance based on the reduction in model accuracy when a given feature is permuted, reported as MeanDecreaseAccuracy. Higher MeanDecreaseAccuracy values indicate genes whose removal most strongly degrades predictive performance, and therefore represent powerful discriminators between experimental conditions.

PLS-DA complements this approach by capturing the contribution of each gene to the latent components that best separate sample classes, quantified by the Variable Importance in Projection (VIP) score. The VIP_mean score reflects how strongly a gene contributes to maximizing class discrimination in reduced-dimensional projection space, with VIP > 1 generally considered biologically meaningful and VIP > 1.3 as highly influential.

By integrating both MeanDecreaseAccuracy and VIP_mean, we prioritized genes that are consistently important across orthogonal modeling frameworks—predictive in a classification context (RF) and structurally informative in discriminant projection space (PLS-DA). This intersection identifies a robust discriminant gene signature representing the molecular drivers underlying the pathway-level rewiring observed in APC knockdown and chemotherapeutic response.

Feature selection using the combined Random Forest and PLS-DA approach yielded a 43-gene discriminant signature representing the strongest molecular contributors to genotype- and treatment-dependent separation (Figure 6 (a)). Genes with the highest MeanDecreaseAccuracy (Random Forest) and VIP_mean (PLS-DA) scores included CCNB3, ORC1, and E2F2, which regulate G2/M transition, replication origin licensing, and E2F-driven transcription, respectively, indicating checkpoint-regulated cell-cycle control and DNA-damage response activation (Figure 6(b)). Additional high-importance genes such as VGF, CXCL2, and IL11 are associated with inflammatory cytokine signaling and stress adaptation, aligning with the NF-κB-mediated survival and cytokine activation observed in Hallmark and Oncogenic pathway analyses. The presence of UNG, a key base-excision repair enzyme, further reinforces the strong DDR signature identified across pathway analyses [54].

**Figure 6(a).**
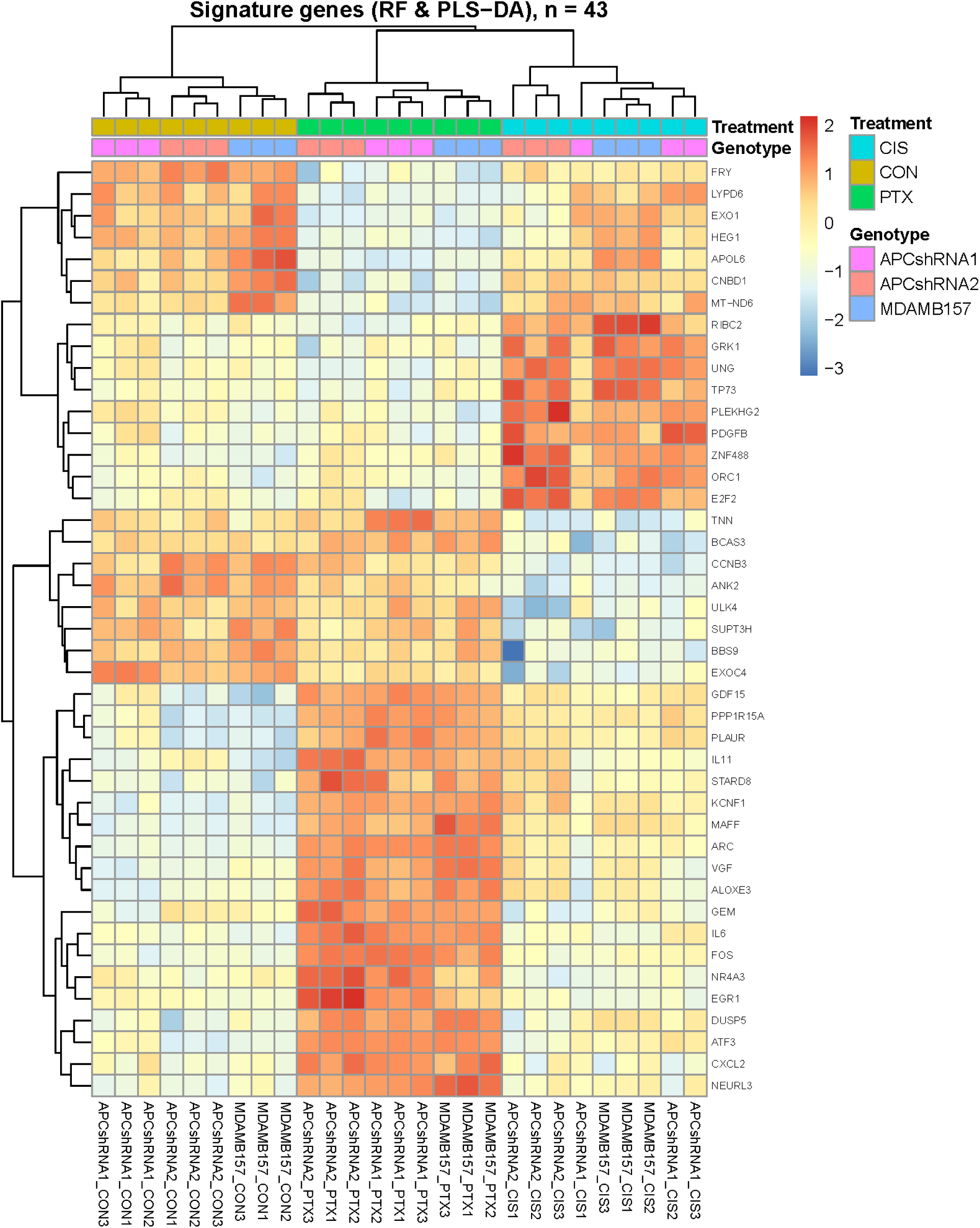
Heatmap of discriminant gene expression signature identified by integrated Random Forest (RF) and PLS-DA analysis (n = 43). Hierarchical clustering of Z-score–scaled expression values show distinct grouping of samples based on genotype (APC_shRNA1, APC_shRNA2, and parental MDA-MB-157) and treatment condition (CON, CIS, PTX). The heatmap highlights a strong block of elevated expression in APC_shRNA1-PTX and APC_shRNA2-CIS groups, consistent with their experimentally validated drug-resistant phenotypes. Key contributors include genes involved in checkpoint regulation (CCNB3, ORC1, E2F2), inflammatory/cytokine signaling (CXCL2, IL11, VGF), and DNA repair (UNG), supporting pathway-level findings of DDR activation, metabolic reinforcement, and inflammatory survival signaling. Overall, these 43 genes represent a robust multi-method discriminant signature capturing genotype- and treatment-dependent transcriptional rewiring.

**Figure 6 (b).**
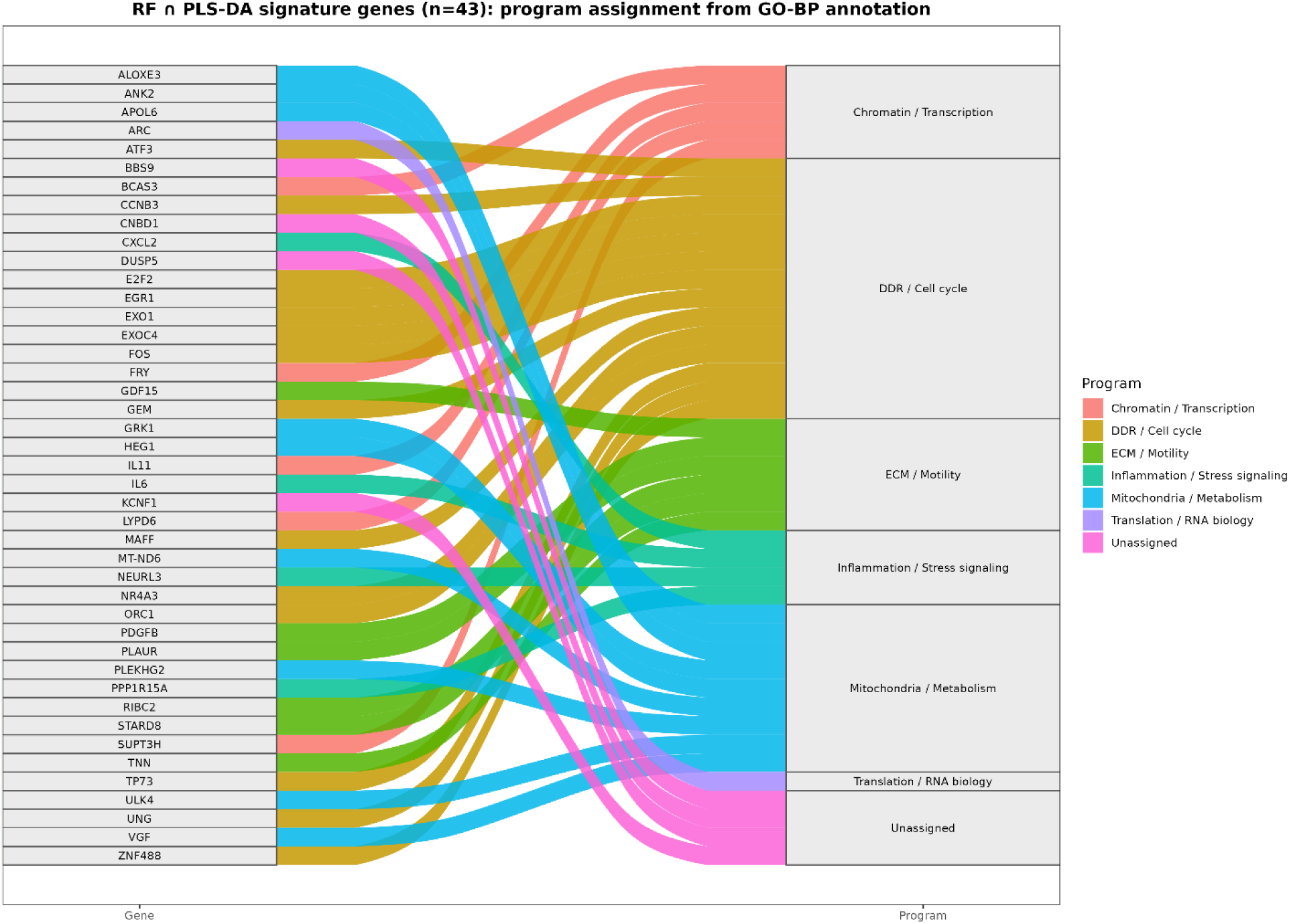
Alluvial plot illustrates the functional convergence of discriminant genes onto shared biological programs. The diagram highlights how genes identified by combined RF and PLS-DA analyses map to major functional categories implicated in DNA damage response, mitochondrial metabolism, inflammatory signaling, and chromatin regulation. Four genes, including *DUSP5*, *BBS9*, *KCNF1*, and *CNBD1*, lack curated GO Biological Process annotations, reflecting limited functional annotation rather than biological irrelevance. These genes were therefore retained in the analysis but could not be assigned to specific GO biological process categories.

Genes linked to signaling and membrane remodeling, including GRK1, KCNF1, TNN, and CNBD1, suggest alterations in ion-channel regulation and membrane signaling architecture, potentially supporting survival under chemotherapy-induced stress. Additionally, mitochondrial and metabolic regulators (e.g., MT-ND6 from the broader list) are consistent with the OXPHOS-intensive metabolic phenotype observed in the enrichment analyses, indicating energy reinforcement to sustain repair and stress-response programs [46].

Together, these findings demonstrate that genes ranked highly by both MeanDecreaseAccuracy and VIP_mean collectively represent a robust multi-method discriminant signature capturing core biological features of APC-driven adaptation, including cell-cycle and DNA-repair control, mitochondrial metabolic support, inflammatory survival signaling, and transcriptional and signaling plasticity. This signature provides a gene-level fingerprint of the broader pathway reprogramming identified in APC knockdown and drug-treated states.

## 4. Conclusion

In this study, we systematically investigated the transcriptomic consequences of APC depletion and chemotherapeutic exposure in the triple-negative breast cancer (TNBC) model MDA-MB-157, revealing extensive remodeling of cellular pathways associated with survival, metabolic adaptation, and stress tolerance. Global dimensionality-reduction analyses (PCA and t-SNE) demonstrated complexity in the separation of samples by genotype, with cisplatin and paclitaxel treatments producing secondary clustering effects that further reflect treatment-specific transcriptional responses. These findings point to biologically meaningful transcriptomic divergence induced by APC knockdown and chemotherapy.

Pathway enrichment analysis integrating Hallmark, GO, and Oncogenic signature gene sets revealed distinct mechanistic architectures underlying APC_shRNA1 (paclitaxel-tolerant) and APC_shRNA2 (cisplatin-tolerant) phenotypes at baseline. APC_shRNA1 cells displayed strong enrichment for EMT, inflammatory NF-κB survival signaling, OXPHOS-dependent metabolic reprogramming, and focal adhesion remodeling, defining a motile, energetically flexible, and anti-apoptotic state consistent with paclitaxel persistence. In contrast, APC_shRNA2 cells demonstrated enhanced antioxidant and redox buffering capacity, DNA-repair preparedness, and extracellular-matrix restructuring, characteristic of a redox-protected, damage-primed cellular state commonly associated with cisplatin tolerance.

In parental MDA-MB-157 cells, cisplatin exposure induced a repair-committed phenotype supported by mitochondrial bioenergetics, checkpoint-regulated cycling, and enhanced proteostasis, while paclitaxel treatment triggered a compensation response involving DNA-damage signaling, OXPHOS reinforcement, detoxification pathways, and chromatin remodeling despite transcriptional suppression of mitotic-spindle structural genes. Together, these profiles reveal that both chemotherapies elicit coordinated adaptation programs that integrate DDR signaling, mitochondrial metabolism, translation, inflammatory circuits, and chromatin accessibility—mechanisms closely linked to persistent minimal-residual-disease behavior.

To identify molecular determinants of these adaptive phenotypes, Random Forest and PLS-DA machine-learning models were applied to derive a robust 43-gene discriminant signature that consistently separated genotypes and treatment states. High-importance markers included cell-cycle regulators (CCNB3, ORC1, E2F2), inflammatory and cytokine mediators (CXCL2, IL11, VGF), DNA-repair enzymes (UNG), and mitochondrial/metabolic components (e.g., MT-ND6), defining a gene-level fingerprint reflecting the DDR-coupled, OXPHOS-supported, transcriptionally plastic survival strategies uncovered by pathway analysis.

Collectively, these results indicate that APC loss drives profound transcriptional rewiring that primes TNBC cells for drug adaptability through interconnected programs involving DNA repair, mitochondrial metabolism, inflammatory signaling, and epigenetic flexibility. Chemotherapeutic exposure amplifies these adaptive systems, enabling checkpoint-governed survival rather than treatment-induced collapse. The RF/PLS-DA-derived signature highlights candidate biomarkers and potential therapeutic vulnerabilities that may guide precision strategies aimed at disrupting the adaptive circuitry underlying treatment resistance in APC-deficient TNBC.

These findings have important translational relevance for the treatment of APC-deficient TNBC. The pathway and machine-learning analyses reveal that APC loss generates a stress-adapted, energetically reinforced, and epigenetically flexible cell state that enables survival under cisplatin and paclitaxel exposure. Rather than undergoing terminal arrest or apoptosis, APC-depleted cells preserve checkpoint-governed proliferation through the coordinated activation of DNA-damage repair networks, mitochondrial oxidative phosphorylation, cap-dependent translation, inflammatory and cytokine signaling, and chromatin remodeling. This survival architecture closely parallels cellular programs associated with chemoresistant minimal residual disease in aggressive breast cancers [52].

The RF + PLS-DA 43-gene discriminant signature highlights actionable vulnerabilities, including DDR regulators (E2F2, ORC1, UNG), inflammatory mediators (CXCL2, IL11), metabolic drivers (MT-ND6), and translation-control effectors (MYC, EIF4E). These results suggest that therapeutic combinations targeting DDR–metabolism–translation axes or disrupting inflammatory/epigenetic plasticity (e.g., OXPHOS inhibitors such as IACS-010759, MYC/translation blockers such as CX-5461, BET/HDAC inhibitors, or NF-κB pathway antagonists) may sensitize APC-deficient tumors and overcome taxane and platinum resistance. The mechanistic clarity provided by this model supports the rational development of biomarker-driven stratification strategies and combination therapies for APC-associated TNBC.

## Acknowledgements

This research was supported by the American Cancer Society – Institutional Research Grant, the Indiana CTSI, grant #UL1 TR001108 from the NIH, NCATS, and a CTSI Core Usage grant (JRP). TMN thanks IUSB faculty research grant and NSF award 1726218 for supporting this research.

## References

[1] F. Bray, J. Ferlay, I. Soerjomataram, R.L. Siegel, L.A. Torre, A. Jemal, Global cancer statistics 2018: GLOBOCAN estimates of incidence and mortality worldwide for 36 cancers in 185 countries, CA Cancer J Clin, (2018).

[2] R.L. Siegel, K.D. Miller, A. Jemal, Cancer statistics, 2018, CA Cancer J Clin, 68 (2018) 7–30.

[3] G. Bianchini, Y. Qi, R.H. Alvarez, T. Iwamoto, C. Coutant, N.K. Ibrahim, V. Valero, M. Cristofanilli, M.C. Green, L. Radvanyi, C. Hatzis, G.N. Hortobagyi, F. Andre, L. Gianni, W.F. Symmans, L. Pusztai, Molecular anatomy of breast cancer stroma and its prognostic value in estrogen receptor-positive and -negative cancers, J Clin Oncol, 28 (2010) 4316–4323.

[4] D.J. Slamon, G.M. Clark, S.G. Wong, W.J. Levin, A. Ullrich, W.L. McGuire, Human breast cancer: correlation of relapse and survival with amplification of the HER-2/neu oncogene, Science, 235 (1987) 177–182.

[5] C.M. Perou, Molecular stratification of triple-negative breast cancers, Oncologist, 16 Suppl 1 (2011) 61–70.

[6] C.M. Perou, Molecular stratification of triple-negative breast cancers, Oncologist, 15 Suppl 5 (2010) 39–48.

[7] T. Sorlie, R. Tibshirani, J. Parker, T. Hastie, J.S. Marron, A. Nobel, S. Deng, H. Johnsen, R. Pesich, S. Geisler, J. Demeter, C.M. Perou, P.E. Lonning, P.O. Brown, A.L. Borresen-Dale, D. Botstein, Repeated observation of breast tumor subtypes in independent gene expression data sets, Proc Natl Acad Sci U S A, 100 (2003) 8418–8423.

[8] I.A. Mayer, V.G. Abramson, B.D. Lehmann, J.A. Pietenpol, New strategies for triple-negative breast cancer--deciphering the heterogeneity, Clin Cancer Res, 20 (2014) 782–790.

[9] K. Chin, S. DeVries, J. Fridlyand, P.T. Spellman, R. Roydasgupta, W.L. Kuo, A. Lapuk, R.M. Neve, Z. Qian, T. Ryder, F. Chen, H. Feiler, T. Tokuyasu, C. Kingsley, S. Dairkee, Z. Meng, K. Chew, D. Pinkel, A. Jain, B.M. Ljung, L. Esserman, D.G. Albertson, F.M. Waldman, J.W. Gray, Genomic and transcriptional aberrations linked to breast cancer pathophysiologies, Cancer Cell, 10 (2006) 529–541.

[10] N. Cancer Genome Atlas, Comprehensive molecular portraits of human breast tumours, Nature, 490 (2012) 61–70.

[11] G. Ciriello, M.L. Gatza, A.H. Beck, M.D. Wilkerson, S.K. Rhie, A. Pastore, H. Zhang, M. McLellan, C. Yau, C. Kandoth, R. Bowlby, H. Shen, S. Hayat, R. Fieldhouse, S.C. Lester, G.M. Tse, R.E. Factor, L.C. Collins, K.H. Allison, Y.Y. Chen, K. Jensen, N.B. Johnson, S. Oesterreich, G.B. Mills, A.D. Cherniack, G. Robertson, C. Benz, C. Sander, P.W. Laird, K.A. Hoadley, T.A. King, T.R. Network, C.M. Perou, Comprehensive Molecular Portraits of Invasive Lobular Breast Cancer, Cell, 163 (2015) 506–519.

[12] C.M. Perou, T. Sorlie, M.B. Eisen, M. van de Rijn, S.S. Jeffrey, C.A. Rees, J.R. Pollack, D.T. Ross, H. Johnsen, L.A. Akslen, O. Fluge, A. Pergamenschikov, C. Williams, S.X. Zhu, P.E. Lonning, A.L. Borresen-Dale, P.O. Brown, D. Botstein, Molecular portraits of human breast tumours, Nature, 406 (2000) 747–752.

[13] A. Prat, C.M. Perou, Deconstructing the molecular portraits of breast cancer, Mol Oncol, 5 (2011) 5–23.

[14] C. Joseph, A. Papadaki, M. Althobiti, M. Alsaleem, M.A. Aleskandarany, E.A. Rakha, Breast cancer intratumour heterogeneity: current status and clinical implications, Histopathology, (2018).

[15] K. Polyak, Heterogeneity in breast cancer, J Clin Invest, 121 (2011) 3786–3788.

[16] D. Zdravkovic, D. Nikolic, M. Zdravkovic, Heterogeneity of tumor cells and metastases in breast cancer patients: cause or consequence?, Breast Cancer Res Treat, (2018).

[17] Y.S. Chang, C.Y. Lin, S.F. Yang, C.M. Ho, J.G. Chang, Analysing the mutational status of adenomatous polyposis coli (APC) gene in breast cancer, Cancer Cell Int, 16 (2016) 23.

[18] C.P. Prasad, S. Mirza, G. Sharma, R. Prashad, S. DattaGupta, G. Rath, R. Ralhan, Epigenetic alterations of CDH1 and APC genes: relationship with activation of Wnt/beta-catenin pathway in invasive ductal carcinoma of breast, Life Sci, 83 (2008) 318–325.

[19] D. Sarrio, G. Moreno-Bueno, D. Hardisson, C. Sanchez-Estevez, M. Guo, J.G. Herman, C. Gamallo, M. Esteller, J. Palacios, Epigenetic and genetic alterations of APC and CDH1 genes in lobular breast cancer: relationships with abnormal E-cadherin and catenin expression and microsatellite instability, Int J Cancer, 106 (2003) 208–215.

[20] I. Van der Auwera, S.J. Van Laere, S.M. Van den Bosch, G.G. Van den Eynden, B.X. Trinh, P.A. van Dam, C.G. Colpaert, M. van Engeland, E.A. Van Marck, P.B. Vermeulen, L.Y. Dirix, Aberrant methylation of the Adenomatous Polyposis Coli (APC) gene promoter is associated with the inflammatory breast cancer phenotype, Br J Cancer, 99 (2008) 1735–1742.

[21] J.R. Prosperi, A.I. Khramtsov, G.F. Khramtsova, K.H. Goss, Apc mutation enhances PyMT-induced mammary tumorigenesis, PLoS One, 6 (2011) e29339.

[22] R. Fodde, J. Kuipers, C. Rosenberg, R. Smits, M. Kielman, C. Gaspar, J.H. van Es, C. Breukel, J. Wiegant, R.H. Giles, H. Clevers, Mutations in the APC tumour suppressor gene cause chromosomal instability, Nat Cell Biol, 3 (2001) 433–438.

[23] K. Sugioka, L.E. Fielmich, K. Mizumoto, B. Bowerman, S. van den Heuvel, A. Kimura, H. Sawa, Tumor suppressor APC is an attenuator of spindle-pulling forces during C. elegans asymmetric cell division, Proc Natl Acad Sci U S A, 115 (2018) E954–E963.

[24] A. Tighe, V.L. Johnson, S.S. Taylor, Truncating APC mutations have dominant effects on proliferation, spindle checkpoint control, survival and chromosome stability, J Cell Sci, 117 (2004) 6339–6353.

[25] J. Groden, G. Joslyn, W. Samowitz, D. Jones, N. Bhattacharyya, L. Spirio, A. Thliveris, M. Robertson, S. Egan, M. Meuth, et al., Response of colon cancer cell lines to the introduction of APC, a colon-specific tumor suppressor gene, Cancer Res, 55 (1995) 1531–1539.

[26] P.J. Morin, B. Vogelstein, K.W. Kinzler, Apoptosis and APC in colorectal tumorigenesis, Proc Natl Acad Sci U S A, 93 (1996) 7950–7954.

[27] C.B. Anderson, K.L. Neufeld, R.L. White, Subcellular distribution of Wnt pathway proteins in normal and neoplastic colon, Proc Natl Acad Sci U S A, 99 (2002) 8683–8688.

[28] T. Fujii, F. Le Du, L. Xiao, T. Kogawa, C.H. Barcenas, R.H. Alvarez, V. Valero, Y. Shen, N.T. Ueno, Effectiveness of an Adjuvant Chemotherapy Regimen for Early-Stage Breast Cancer: A Systematic Review and Network Meta-analysis, JAMA Oncol, 1 (2015) 1311–1318.

[29] R. Bartsch, E. Bergen, A. Galid, Current concepts and future directions in neoadjuvant chemotherapy of breast cancer, Memo, 11 (2018) 199–203.

[30] Z. Nahleh, Neoadjuvant chemotherapy for “triple negative” breast cancer: a review of current practice and future outlook, Med Oncol, 27 (2010) 531–539.

[31] J. Schwartz, Current combination chemotherapy regimens for metastatic breast cancer, Am J Health Syst Pharm, 66 (2009) S3–8.

[32] P.E.R. Spronk, K.M. de Ligt, A.C.M. van Bommel, S. Siesling, C.H. Smorenburg, M. Vrancken Peeters, N.B.C. Audit, Current decisions on neoadjuvant chemotherapy for early breast cancer: Experts’ experiences in the Netherlands, Patient Educ Couns, (2018).

[33] M.M. Gottesman, Mechanisms of cancer drug resistance, Annu Rev Med, 53 (2002) 615–627.

[34] S.M. Bevill, J.S. Zawistowski, G.L. Johnson, Enhancer remodeling regulates epigenetic adaptation and resistance to MEK1/2 inhibition in triple-negative breast cancer, Mol Cell Oncol, 4 (2017) e1300622.

[35] G. Greville, A. McCann, P.M. Rudd, R. Saldova, Epigenetic regulation of glycosylation and the impact on chemo-resistance in breast and ovarian cancer, Epigenetics, 11 (2016) 845–857.

[36] R. Hubaux, K.L. Thu, W.L. Lam, Re: the Wnt signaling pathway in non-small cell lung cancer, J Natl Cancer Inst, 106 (2014).

[37] D.J. Stewart, Wnt signaling pathway in non-small cell lung cancer, J Natl Cancer Inst, 106 (2014) djt356.

[38] M.K. VanKlompenberg, C.O. Bedalov, K.F. Soto, J.R. Prosperi, APC selectively mediates response to chemotherapeutic agents in breast cancer, BMC Cancer, 15 (2015) 457.

[39] M.K. VanKlompenberg, E. Leyden, A.H. Arnason, J.T. Zhang, C.D. Stefanski, J.R. Prosperi, APC loss in breast cancer leads to doxorubicin resistance via STAT3 activation, Oncotarget, 8 (2017) 102868–102879.

[40] F. Koopmans, GOAT: efficient and robust identification of gene set enrichment, Commun Biol, 7 (2024) 744.

[41] L. Breiman, Random Forests, Machine Learning, 45 (2001) 5–32.

[42] S. Wold, M. Sjöström, L. Eriksson, PLS-regression: a basic tool of chemometrics, Chemometrics and Intelligent Laboratory Systems, 58 (2001) 109–130.

[43] F. Rohart, B. Gautier, A. Singh, K.A. Le Cao, mixOmics: An R package for ‘omics feature selection and multiple data integration, PLoS Comput Biol, 13 (2017) e1005752.

[44] Y. Wang, C. Dong, B.P. Zhou, Metabolic reprogram associated with epithelial-mesenchymal transition in tumor progression and metastasis, Genes Dis, 7 (2020) 172–184.

[45] A. Datta, S. Deng, V. Gopal, K.C. Yap, C.E. Halim, M.L. Lye, M.S. Ong, T.Z. Tan, G. Sethi, S.C. Hooi, A.P. Kumar, C.T. Yap, Cytoskeletal Dynamics in Epithelial-Mesenchymal Transition: Insights into Therapeutic Targets for Cancer Metastasis, Cancers (Basel), 13 (2021).

[46] S. Bao, X. Wang, M. Li, Z. Gao, D. Zheng, D. Shen, L. Liu, Potential of Mitochondrial Ribosomal Genes as Cancer Biomarkers Demonstrated by Bioinformatics Results, Frontiers in Oncology, Volume 12 - 2022 (2022).

[47] E.C. Koc, F.C. Koc, F. Kartal, M. Tirona, H. Koc, Role of mitochondrial translation in remodeling of energy metabolism in ER/PR(+) breast cancer, Frontiers in Oncology, Volume 12 - 2022 (2022).

[48] M. Devanaboyina, J. Kaur, E. Whiteley, L. Lin, K. Einloth, S. Morand, L. Stanbery, D. Hamouda, J. Nemunaitis, NF-kappaB Signaling in Tumor Pathways Focusing on Breast and Ovarian Cancer, Oncol Rev, 16 (2022) 10568.

[49] G. Barrera, M.A. Cucci, M. Grattarola, C. Dianzani, G. Muzio, S. Pizzimenti, Control of Oxidative Stress in Cancer Chemoresistance: Spotlight on Nrf2 Role, Antioxidants (Basel), 10 (2021).

[50] L. Gan, Y. Yang, Q. Li, Y. Feng, T. Liu, W. Guo, Epigenetic regulation of cancer progression by EZH2: from biological insights to therapeutic potential, Biomark Res, 6 (2018) 10.

[51] W. Jiang, X. Wang, C. Zhang, L. Xue, L. Yang, Expression and clinical significance of MAPK and EGFR in triple-negative breast cancer, Oncol Lett, 19 (2020) 1842–1848.

[52] C.J. Lord, A. Ashworth, The DNA damage response and cancer therapy, Nature, 481 (2012) 287–294.

[53] J. Gallego-Jara, G. Lozano-Terol, R.A. Sola-Martinez, M. Canovas-Diaz, T. de Diego Puente, A Compressive Review about Taxol((R)): History and Future Challenges, Molecules, 25 (2020).

[54] L.D. Weeks, G.E. Zentner, P.C. Scacheri, S.L. Gerson, Uracil DNA glycosylase (UNG) loss enhances DNA double strand break formation in human cancer cells exposed to pemetrexed, Cell Death Dis, 5 (2014) e1045.

